# Early cell cycle genes in cortical organoid progenitors predict interindividual variability in infant brain growth trajectories

**DOI:** 10.1101/2025.02.07.637106

**Authors:** Madison R. Glass, Nana Matoba, Alvaro A. Beltran, Niyanta K. Patel, Tala M. Farah, Karthik Eswar, Shivam Bhargava, Karen Huang, Ian Curtin, Sara Ahmed, Mary Srivastava, Emma Drake, Liam T. Davis, Meghana Yeturi, Kexin Sun, Michael I. Love, Jeremy M. Simon, Tanya St. John, Natasha Marrus, Juhi Pandey, Annette Estes, Stephen Dager, Robert T. Schultz, Kelly Botteron, Alan Evans, Sun Hyung Kim, Martin Styner, Robert C. McKinstry, D. Louis Collins, Heather Volk, Kelly Benke, Lonnie Zwaigenbaum, Heather Hazlett, Adriana S. Beltran, Jessica B. Girault, Mark D. Shen, Joseph Piven, Jason L. Stein, the Infant Brain Imaging Study Network

**Affiliations:** Department of Genetics, University of North Carolina at Chapel Hill; Chapel Hill, NC, USA; UNC Neuroscience Center, University of North Carolina at Chapel Hill; Chapel Hill, NC, USA; Department of Biostatistics, University of North Carolina at Chapel Hill; Chapel Hill, NC, USA; Department of Speech and Hearing Sciences, University of Washington; Seattle, WA, USA; University of Washington Autism Center, University of Washington; Seattle, WA, USA; Department of Psychiatry, Washington University School of Medicine; St. Louis, MO, USA; Department of Psychiatry, University of Pennsylvania; Philadelphia, PA, USA; Center for Autism Research, Children’s Hospital of Philadelphia; Philadelphia, PA, USA; Department of Radiology, University of Washington; Seattle, WA, USA; Department of Bioengineering, University of Washington; Seattle, WA, USA; Department of Pediatrics, University of Pennsylvania; Philadelphia, USA; Department of Neurology and Neurosurgery, McGill University; Montreal, QC, CA; Department of Psychiatry, McGill University; Montreal, QC, CA; Department of Biomedical Engineering, McGill University; Montreal, QC, CA; Carolina Institute for Developmental Disabilities, University of North Carolina at Chapel Hill; Chapel Hill, NC, USA; Department of Psychiatry, University of North Carolina at Chapel Hill; Chapel Hill, NC, USA; Department of Computer Sciences, University of North Carolina at Chapel Hill; Chapel Hill, NC, USA; School of Public Health, Johns Hopkins; Baltimore, MD, USA; Department of Developmental Pediatrics, University of Alberta; Edmonton, AB, CA

**Keywords:** Cell cycle, neurogenesis, cortical hem, WNT, organoid, infant brain growth

## Abstract

Human induced pluripotent stem cell (iPSC) derived cortical organoids (hCOs) model neurogenesis on an individual’s genetic background. The degree to which hCO phenotypes recapitulate the brain growth of the participants from which they were derived is not well established. We generated up to 3 iPSC clones from each of 18 participants in the Infant Brain Imaging Study, who have undergone longitudinal brain imaging during infancy. We identified consistent hCO morphology and cortical cell types across clones from the same participant. hCO cross-sectional area and production of cortical hem cells were associated with *in vivo* cortical growth rates. Cell cycle associated genes expression in early progenitors at the crux of fate decision trajectories were correlated with cortical growth rate from 6-12 months of age, and were enriched in microcephaly and neurodevelopmental disorder genes. Our data suggest the hCOs capture inter-individual variation in cortical cell types influencing infant cortical surface area expansion.

## Introduction

Induced pluripotent stem cells (iPSCs) differentiated into human cortical organoids (hCOs) contain cortical neural progenitors and their neuronal progeny harboring the same genetic variants as the individual from which they were derived. hCOs have been shown to mimic the developing human cortex through both transcriptomic similarities to fetal tissue and 3D organization in neuroepithelial buds that resemble the developing cortical wall^1–3^. Cortical organoid differentiation generates consistent proportions of cell types and structure both within and across labs^4–6^. However, hCOs have high metabolic stress, imperfect fate specification, and model only a subset of cell types that exist in the *in vivo* brain^3,7^. Given the limitations of current hCOs, the fidelity, or degree to which hCOs recapitulate the inter-individual variability of brain development in individuals from which they were derived, is largely unknown.

Previous studies have used hCOs to study inter-individual variation, though in a limited number of participants or without reference to the *in vivo* phenotypes of the individuals from which they were derived^8–11^. hCOs mirror brain size changes when harboring microcephaly or macrocephaly associated mutations, supporting their fidelity for modeling brain size^12,13^. In addition, cellular and molecular characteristics of two-dimensional neural cultures and hCOs are associated with head or brain size measurements from individuals with autism spectrum disorder (ASD) in the absence of known causal single gene variants^14–17^. However, assessing the fidelity of hCOs for modeling inter-individual variability in cortical size has been limited in previous studies by low sample size, cross-sectional measurements of *in vivo* cortical structure, and lack of early postnatal developmental trajectories starting from infancy.

The fidelity of hCOs to model cortical structure is relevant to ASD, as individuals with high familial likelihood have abnormally increased cortical surface area growth rates during infancy, prior to diagnosis, leading to increased brain size^18–20^. One well-known cellular mechanism that results in increased cortical surface area across species is the expansion of the neural progenitor pool present during fetal development^21^. Increased neural progenitor proliferation rate, increased self-renewal of progenitors in early development, or an increased production of transiently amplifying progenitor cells such as outer radial glia (oRG) or intermediate progenitors (IP) result in an expanded neural progenitor pool^22^. An increase in the total number of neural progenitors leads to an increase in the amount of neurons produced and consequently larger cortical surface area.

To assess the fidelity of hCOs for modeling cortical structure, we generated iPSCs from 18 participants in the Infant Brain Imaging Study (IBIS) who underwent at least two longitudinal brain magnetic resonance imaging (MRI) scans and extensive behavioral and clinical testing throughout infancy (**Figure 1a**)^18^. Up to three clones were generated per participant to assess reproducibility. We leveraged this unique, deeply phenotyped set of participants to examine the relationship between organoid growth, cell type production, and gene expression at a signal cell resolution with longitudinal cortical surface area measurements, finding that organoid size, cell type proportions, and progenitors undergoing fate decisions are associated with inter-individual differences in cortical surface area growth rates. Our results demonstrate the degree of reproducibility and fidelity of hCOs, while also identifying cell types and genes underlying inter-individual differences in human cortical structure.

**Fig. 1.**
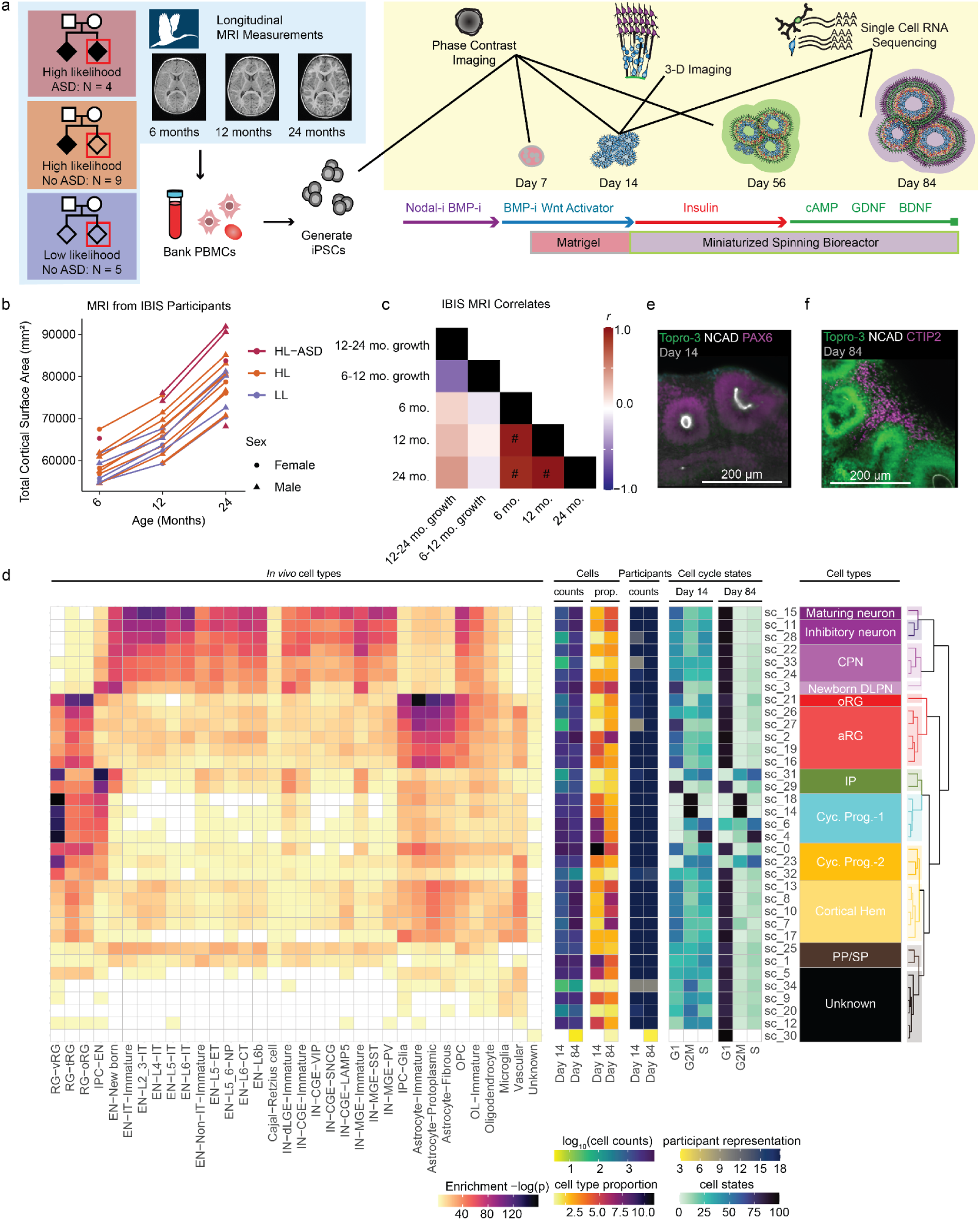
Strategy to Model Inter-Individual Variation in Infant MRI Measures using Cortical Organoids. **a**, Experimental Overview. Previous Infant Brain Imaging Study (IBIS) research collected longitudinal MRI measurements from three groups of participants: Infants with no family history (low-likelihood, LL) of neuropsychiatric disorders, infants with an older sibling diagnosed with autism spectrum disorder (ASD) who either were not diagnosed at 24 months of age (high likelihood, HL) or received an ASD diagnosis (high likelihood - ASD, HL-ASD). Up to 3 iPSC clones were generated from previous IBIS participants’ PBMCs. iPSC lines were differentiated into cortical organoids and assayed with scRNAseq at day 14 and day 84, 3D imaged at day 14, and imaged throughout the differentiation to measure cross-sectional area and organoid morphology. **b,** MRI measurements of cortical surface area in IBIS participants in the study. **c**, correlations among MRI measurements of IBIS participants in this study. # indicates FDR < 0.1 **d,** scRNAseq cell type annotations. **e,** Representative day 14 organoid showing NCAD+ neuroepithelial buds with PAX6+ radial glia. **f,** Day 84 organoids showing CTIP2+ lower layer neurons outside of neuroepithelial buds. Abbreviation: oligodendrocytes (OL), oligodendrocyte progenitor cells (OPC), intermediate progenitor cell (IPC), excitatory neuron (EN), inhibitory neuron (IN), radial glia (RG), apical radial glia (aRG), outer radial glia (oRG), truncated radial glia (tRG), ventral radial glia (vRG), cortical projection neuron (CPN), deep layer projection neuron (DLPN), Cycling progenitor (Cyc. prog.), pre-plate (PP), subplate (SP).

## Results

### IBIS-iPSC generation, validation, and differentiation

IBIS is a prospective study of infant siblings which seeks to identify behavioral or brain imaging traits during infancy that may predict an ASD diagnosis by two years of age^22,23^. Participants were either from a family with no history of ASD (low likelihood; LL) or had an older sibling who had previously been diagnosed with ASD, leading to an increased likelihood of the infant sibling receiving an ASD diagnosis (high likelihood; HL) (**Figure 1a**)^24^. We recruited 18 total participants including those from HL families who were diagnosed with ASD at 24 months (N = 4), participants from HL families who were not diagnosed with ASD (N = 9), and those from LL families who were not diagnosed with ASD (N = 5). IBIS participants enrolled in this study had at least two MRI-derived measurements of cortical surface area (N = 13 at 6 months, N = 16 at 12 months, N = 15 at 24 months), and contiguous absolute growth rates were calculated when data was available (N = 12 for 6-12 month growth rates, N = 13 for 12-24 month growth rate; **Figure 1b**). Cortical surface area increased from 6 months of age to 24 months of age with expected variability across participants (s.d. = 7,070 mm^2^ at 24 months). Cross-sectional cortical surface area measurements were highly correlated with each other, indicating those with larger brains earlier in development retain larger brains later in development (false discovery rate using Benjamini-Hochberg method (FDR) < 0.1, **Figure 1c**), similar to previous descriptions^25^. In contrast, longitudinal growth rates were not significantly correlated with each other or cross-sectional measurements, suggesting these measurements are independent assessments of brain growth. Previous studies of infant MRI have revealed faster growth during the first year of life, consistent with the data in this study^25,26^.

iPSCs from each participant were reprogrammed from expanded erythroblasts using standard protocols in batches containing multiple diagnostic categories to prevent confounding^27^. Up to three clones per participant were generated to evaluate reproducibility. All clones passed quality control including: Taqman scorecard to assess pluripotency, competence for tri-lineage differentiation, and clearance of Sendai virus used during reprogramming^28^; immunofluorescence for canonical stem cell markers (TRA-180, SOX2, OCT4, SSEA4); and genotyping microarrays to both verify participant identity and check for any reprogramming induced aneuploidy^29^ (**Figure S1**).

IBIS-iPSC clones were differentiated to hCOs blinded to diagnostic status, family history, and cortical surface area (54 total differentiations from 18 participants with 1-3 clones per participant) (**Table S1**). We followed a miniaturized spinning bioreactor protocol adapted to feeder-free conditions that has previously yielded reproducible cell type composition and cortical wall-like structures across multiple sites^4,30,31^. Across the differentiation, we evaluated multiple phenotypes: cell type composition and gene expression were assessed at day 14 (to capture early progenitors) and day 84 (to capture neuronal production) via single-cell RNA-seq (scRNA-seq)^32^; tight junctions (N-cadherin) and radial glia (PAX6) were imaged using whole hCO immunolabeling via tissue clearing at day 14; and hCO morphology and growth via phase contrast imaging were acquired throughout differentiation. This multi-modal design coupled from a well characterized cohort of infants enabled the identification of hCO phenotypes that relate to infant cortical surface area.

### hCOs Produce Cortical Progenitors and Neurons

We used scRNA-seq to identify cell types present within hCOs and to assess their similarity to primary cortical tissue. scRNA-seq data underwent standard pre-processing to remove low-quality or stressed cells, resulting in 210,016 cells for analysis (**Figure S2**)^33–35^. Cells were grouped into 35 subclusters based on transcriptomic similarity, aggregated into major cell classes through hierarchical clustering, and annotated based on expression of marker genes and comparison with primary fetal tissue (**Figures 1d, S3, Table S2**)^36^. Progenitor populations were further classified as cycling progenitors (Cyc. Prog.) by expression of cell cycle genes. Cell classes included expected cell types from neocortical differentiation^17,37^, including apical radial glia (aRG; *SOX2*+, *PAX6*+), outer radial glia (oRG; *PTPRZ1*+, *FOXG1*+), intermediate progenitors (IP; *EOMES*+), the earliest born preplate/subplate neurons (PP/SP; *SNCA*+^38^, *CCN2*+^39^, *TBR1*+), cortical projection neurons (CPN; *BCL11B*+, *TBR1*+), newborn deep layer projection neurons (Newborn DLPN; *EOMES*+, *TBR1*+), upper layer neurons (ULN; *CUX2*+, *SLC17A6*+), cortical hem (hem; *TTR*+, *EMX2*+), and inhibitory neurons (IN; *GAD2*+, *LHX5*+). We observed impaired subtype specification, where hCO cell types match with multiple *in vivo* cell types, as has been previously documented^7^. Immunolabeling of intact hCOs confirmed the cell types identified through scRNA-seq, where N-cadherin lined the lumen of RG dense (PAX6+) neuroepithelial buds at day 14 (**Figure 1e**), EOMES+ IPs and HOPX+ oRGs were present at day 56 (**Figures S3f,g)**, and CTIP2+ and CUX1+ excitatory neurons were present at day 84 (**Figures 1f, S3h**). Cyst structures within hCOs at day 84 were TTR+, a marker for cortical hem (**Figure S3i**).

Unknown cells (12.5%) did not match well with primary tissue and are likely the result of off-target differentiations and were a majority of the metabolically stressed cells (55.8%). Cycling progenitors at day 14 were positively correlated with unknown cell types at day 84, whereas aRGs at day 14 were negatively correlated with unknown cell types at day 84, demonstrating that progenitor behavior early in differentiation is associated with cell types present later in differentiation (**Figures S4a-c**). Unknown cells were retained for identification of technical variables that influence differentiation outcomes, but were excluded from analyses correlating with cortical surface area measurements. Together, these data suggest the hCO differentiation generated expected cortical progenitors and neuronal cell types, allowing the assessment of hCO phenotypes that relate to inter-individual variability in cortical surface area.

### Reproducibility of hCO differentiation across clones from the same participant

To assess the consistency of hCO differentiation across multiple iPSC clones from the same participant, we first assessed organoid morphology through phase contrast imaging at day 14 and 56 (**Figures 2a,b**). Neuroepithelial buds are an indicator of successful cortical differentiation, whereas extension into the matrigel is an indicator of off-target differentiation at day 14^40,41^. Transparent cyst-like outgrowths were also seen as indicators of non-neuronal cell types, likely choroid plexus, that were readily identifiable at day 56^4,40,42^. Trained raters categorized all hCO as having neuroepithelial buds and/or growth into matrigel at day 14, and as having cysts at day 56 using binary classifications of phase contrast images (**Figure 2c**). hCO differentiations from multiple clones of the same participant had more similar morphology scores than organoids from different participants, suggesting that genetic background influences the gross morphology of iPSC-derived hCOs (**Figure 2d**).

**Fig. 2.**
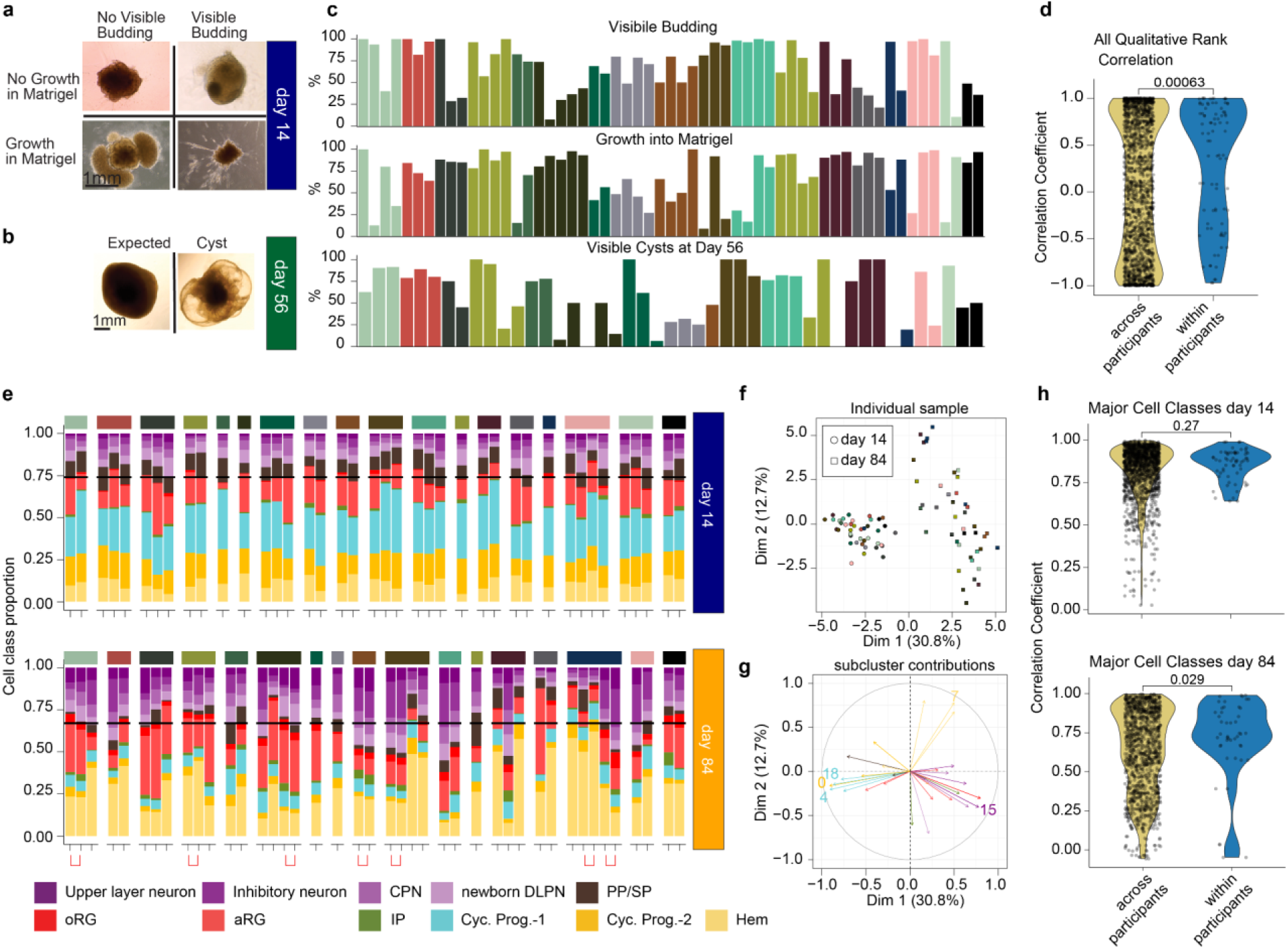
Consistency of differentiation across participants in morphology and cell class proportions. **a**, Day 14 morphology rank indicates if visible neuroepithelial buds were present or if there was growth into the matrigel. **b,** Day 56 morphology rank indicates if clear cysts were present on the organoid. **c**, Quantification of percent rank across day 14 and day 56 for each clone. **d,** Correlation across and within participants for all percent morphological ranks. **e,** Cell class proportion for each library. Libraries from the same differentiation day and clone prepared in different scRNA-seq library preparations are indicated by a bracket. **f**, PCA plot of cell class proportions for samples passing QC. **g**, Cell type proportions contributing to principal components. The top 5 contributing subclusters are labeled. **h**, Correlation of major cell classes across and within participants at day 14 (top) and day 84 (bottom).

Given the variability in morphology across hCOs, we expected that some hCOs underwent unexpected differentiations. hCOs were randomly selected from each clone’s differentiation for scRNA-seq at day 14 whereas smaller hCOs without cysts were selected for scRNA-seq at day 84. Even with this selection criteria, we found that some hCOs were outliers in cell type proportions, likely indicating unexpected differentiations. We excluded scRNA-seq samples where the first principal component of either cell type proportion or gene expression did not cluster with the collection time point or where the proportions of neurons was greater than 2 standard deviations from the average (**Figure S5**). Because the experimental design utilized multiple clones per participant, 18 participants were represented at day 14 (after excluding 4 samples), and 17 participants were represented at day 84 (after excluding 12 samples) following these quality control measures.

After quality control, we found a small increase in neuron proportions (25% to 30%) from day 14 to day 84 and a subsequent decrease in progenitor cells, as expected during neurogenesis (**Figure 2e**). Principal component analysis of cell type proportions shows clear clustering by day of collection (as expected given our quality control criteria), with more variability observed at day 84 after longer differentiations (**Figure 2f**). Variability in cell type proportions from the principal component analysis (PCA) was largely driven by differences in dividing progenitors (weighted toward day 14) and neuronal (weighted more to day 84) subcluster proportions (**Figure 2g**). At day 14, we observed highly consistent cell type proportions across all clones and all participants. Likely because of this global consistency, we did not detect significant differences in cell type proportions within participants as compared to across participants (**Figure 2h**). However, at day 84 when more variability in cell type proportions was present, we did find that cell types were more similar within clones of the same participant as compared to clones across participants. Overall, this suggests that hCO are reliably capturing inter individual variation during the process of neurogenesis.

### Technical Variables Associated with Differentiation Outcomes

While genetic variation contributes to hCO differentiation as demonstrated above, we also tested the degree to which technical variables during iPSC reprogramming, iPSC state, and differentiation handling may also contribute to variability^4,28,43,44^. Though undifferentiated cell state, as marked by individual gene expression in the Taqman scorecard, significantly correlated to some cell type proportions after differentiation, no consistent associations across multiple pluripotency or germ layer markers was observed (**Figure S4d**), likely because all clones were selected to be pluripotent. The Taqman scorecard does not assess if iPSCs were in a naive or primed cell state^45,46^, so we additionally assessed qPCR markers of these states in a subset of the differentiations (12/54). Both naive and primed markers at pluripotency were associated with the percentage of PP/SP neurons after 84 days of differentiation, suggesting that iPSC cell state is pertinent to hCO cell type composition (**Figure S4e**). Though technical factors during reprogramming and differentiation were also correlated to cell type proportions, participants were randomly assigned to rounds of reprogramming or differentiation which should decrease confounding effects due to these technical variables.

Organoid morphology has been associated with cell type composition^40^, and is frequently included as a quality control assessment in organoid differentiation methods^30,41,47^. We identified correlations between hCO cross-sectional area, morphology, NCAD+/PAX6, NCAD+/volume, PAX6+/volume and total volume from intact organoids with cell type proportions (**Figure S6a-d**). For example, visible budding at day 14 was associated with larger hCOs at day 0, 7 and 14, increased proportions of Cyc. Prog.-2 and cysts, and was negatively associated with ULN proportions at day 14 (**Figure S6**), suggesting the visible budding is associated with more progenitor proliferation and less neurogenesis. Cysts at day 56 were also associated with increased unknown cell type proportions at day 14 and increased day 0-7 growth rates (**Figure S6b,c**). Increased proliferation within an hCO may lead to increased cell stress and consequently unknown cell type composition at day 84 (**Figure S4d**). Measurements from intact immunolabeled hCOs did not show correlation to cross-sectional area or rank measurements, or participants’ cortical surface area measurements, possibly due to tissue shrinkage in tissue clearing protocols (**Figure S6e,f**). Our data support using hCO morphology and growth rate may be useful to understand aspects of cellular composition.

### Cell Type Proportion Correlates to Cortical Surface Area and Growth Rate

To examine the relationship between hCOs and infant cortical surface area, we correlated cell type proportions to MRI measurements for each participant (**Figure 3a,b**; **Table S4**). Because of known sex differences in infant cortical surface area, we statistically controlled for the effect of sex^26^. The proportion of day 14 cortical hem cells, as both a major cell class and subcluster (sc_8), positively correlated to 12-24 mo. cortical surface area growth rate. A different subcluster of cortical hem (sc_10) at day 14 was associated with increased 6 month cortical surface area. During brain development, the cortical hem secretes WNT, which regulates cortical patterning and radial glia proliferation^48,49^. Day 84 IP proportions negatively correlated with 6-12 month cortical surface area growth rate, driven by one clone of one participant. Several relationships between the proportion of cortical hem subclusters and major cell class proportions were identified (**Figures 3c-e**). Common genetic variants responsive to WNT signaling have been associated with inter-individual variants in cortical surface area^50^. These relationships suggest that increased endogenous WNT signaling from more cortical hem cells early in development lead to increased progenitor proliferation, neuronal production and ultimately increase cortical surface area during infancy.

**Fig. 3.**
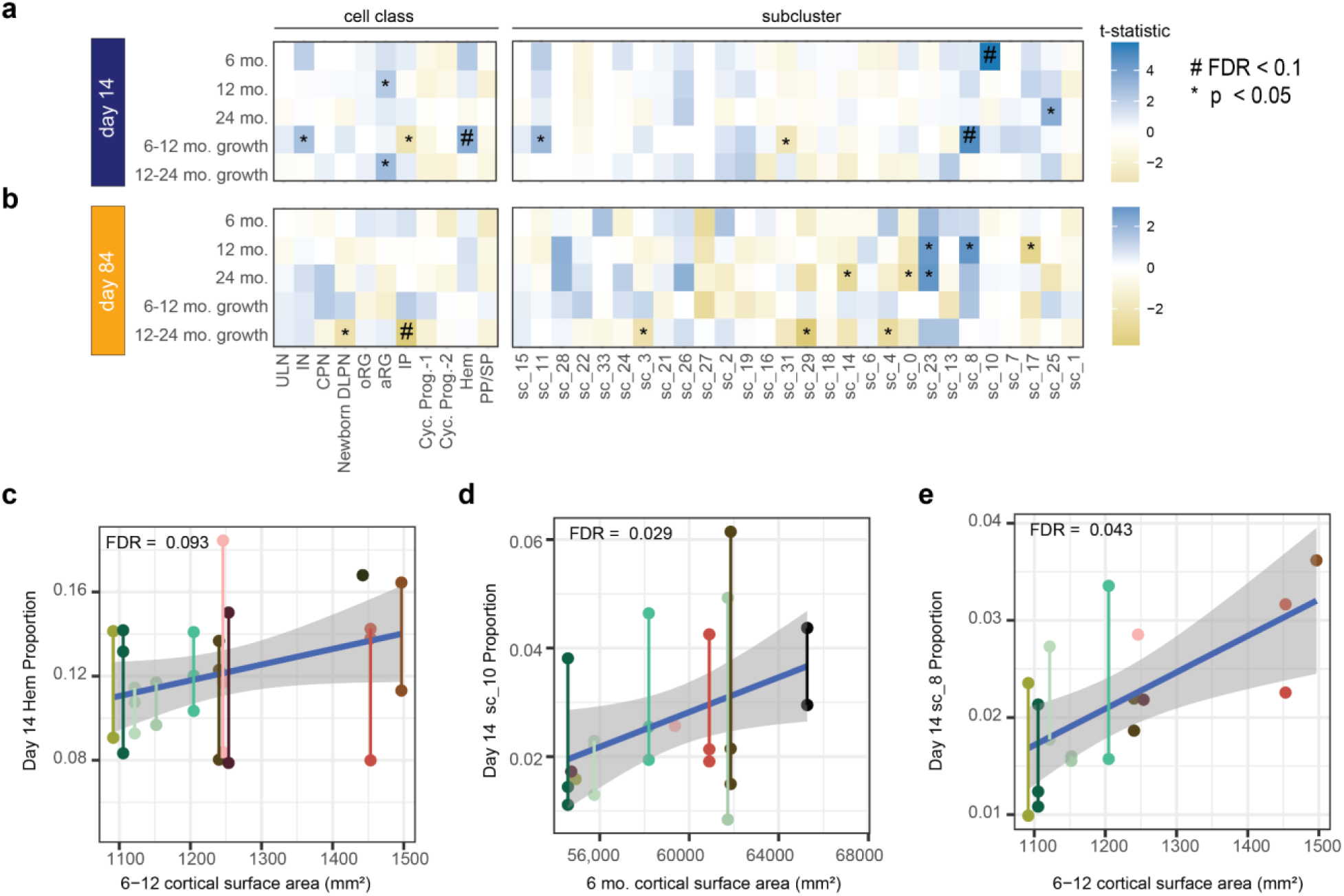
Cell Type Proportion Correlate to MRI. **a**. Average day 14 cell type proportions associated with cortical surface area measurements. **b**. Average day 84 cell type proportions associated with cortical surface area measurements. FDR was calculated for either subclusters or cell classes at each timepoint across MRI measurements. Correlation between cortical surface area growth between 6-12 months of age and day 14 for **c**. cortical hem **d.** sc_10 (cortical hem) and **e.** sc_8 (cortical hem) proportions. Sex is added as a covariate and age at MRI is a covariate for cross-sectional MRI measurements. Day 84 IP association with 12-24 month growth rate was driven by one clone of one donor, so is not highlighted as a scatterplot.

### Cross-sectional Area and 3D Imaging Correlate to Cell Type Proportions

Microcephaly and macrocephaly associated variants affect hCO size, demonstrating that genetics changing progenitor behavior in the developing brain have similar effects *in vitro^12,41^*. We hypothesized that organoid size measured by cross-sectional area changes and intact imaging could also capture inter-individual variation in cortical surface area measurements without harboring variants of strong effect. Cross-sectional organoid area measurements were quantified from phase contrast images taken at day 0, 7, 14 and 35 (**Figure 4a**). A few differentiations shrunk in the Matrigel embedding, likely due to cell migration into the matrix. Smaller organoids at the start of differentiation were associated with slower growth rates, whereas growth rates after 7 days were associated with increased expansion. Average cross sectional hCO area was more similar within than across participants, demonstrating reproducibility (**Figure 4b**).

**Fig. 4.**
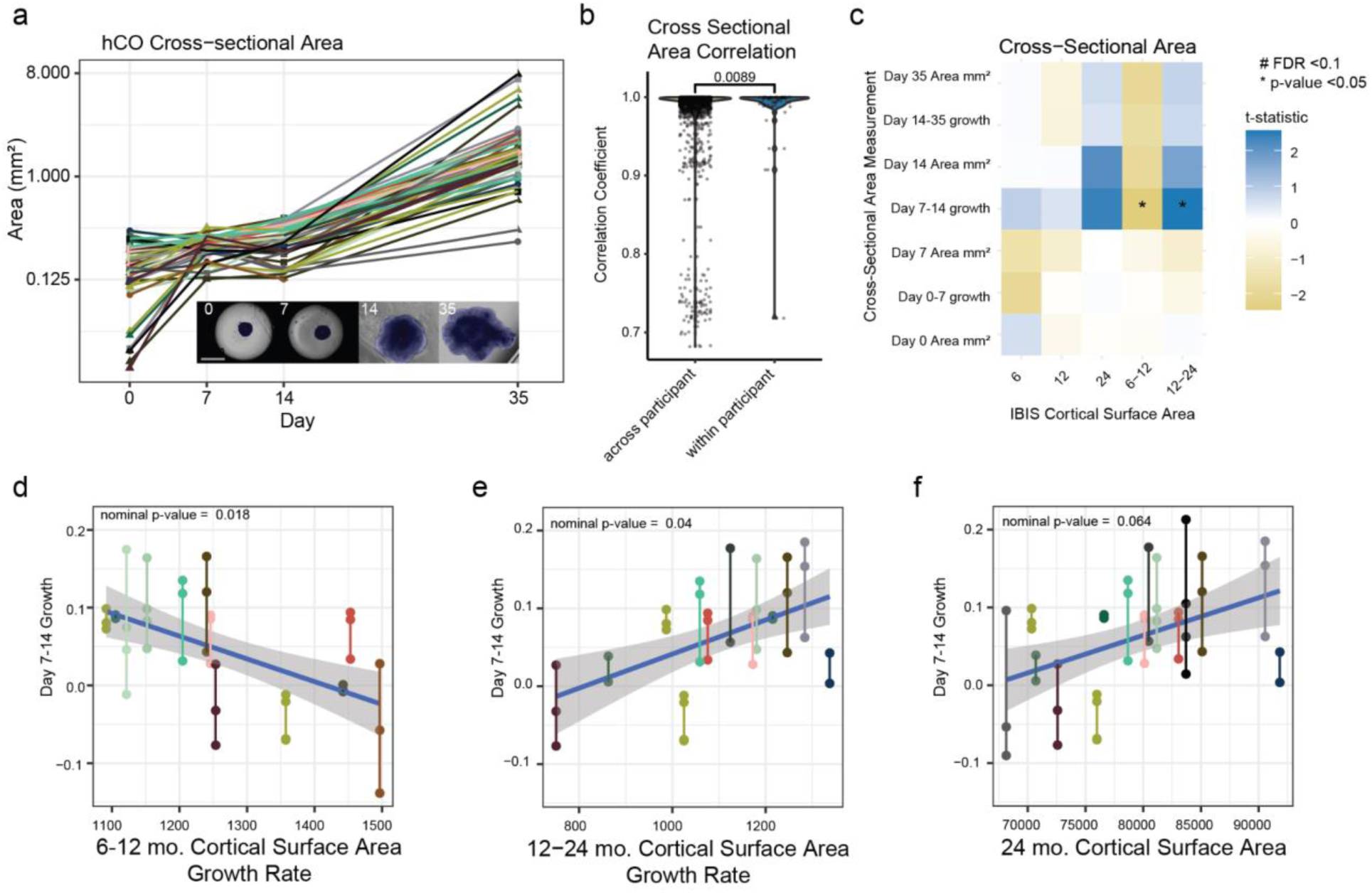
hCO Cross Sectional associations with cortical growth. **a.** hCO cross-sectional area growth rate for each differentiation over time and example mask of an area measurement across differentiation days (inset). **b.** Correlation of cross-sectional area measurements at all timepoints across and within participants. **c.** Cross-sectional area correlations to cortical structure measurements using the average measurements across all differentiated clones. FDR correction was applied for each area measurement to all MRI measurements. Correlations between day 7 and day 14 hCO growth rate with **d.** 6-12 month cortical surface area growth rate, **e.** 12-24 month cortical surface area growth rate and **f.** 24 month cortical surface area.

We then tested if hCO cross-sectional area or growth rate measurements correlated with cortical surface area growth trajectories. The growth rate of hCOs between day 7 and day 14, when organoids were embedded in Matrigel and mostly progenitor cells, nominally correlated with cortical surface area growth rates (**Figure 4c**). hCO growth during the second week of differentiation was negatively associated with absolute cortical surface area growth rate from 6-12 months (**Figure 4d**), and was positively correlated with 12-24 month absolute change in cortical surface area (**Figure 4e**). The opposite direction is consistent with the anti-correlation between the 6-12 mo. and 12-24 mo. growth rate measurements, which potentially measure biologically distinct processes (**Figure 1b**). Though none of the cross-sectional cortical surface area measurements were significant following FDR correction, the direction of the correlation is the same, as would be expected for highly correlated measurements (**Figure 4f**). While differences in hCO seeding likely influenced hCO size later in differentiation as indicated by association with iPSC thaw date, our results suggest that organoid growth rates, which likely normalize for seeding differences, rather than cross-sectional area at a specific time point showed the strongest relationship with brain growth trajectories (**Figure S6b-d**). Other research groups have identified hCO size correlates to macrocephaly and microcephaly^41,51^, and correlations to ADOS severity scores^16^, supporting a link between cross-sectional hCO area as an output of progenitor self-renewal *in vitro* and progenitor pool expansion increasing brain size *in vivo^21^*. Our results suggest that organoid size is related to the cortical surface area of the participant from which it was derived.

### Relationship of gene expression and MRI measurements

We next examined the relationship between gene expression within cell-types and *in vivo* cortical surface area measurements. We focused on sub-clusters of gene expression rather than cell classes to gain greater specificity. We identified 5,161 unique genes significantly correlated to at least one *in vivo* cortical surface area measurement (**Figure 5, Table S6**). Many of the cortical surface area associated genes were found in sc_22 (a cluster within CPN) at day 14. Specifically, 3,087 genes were correlated to growth rate from 6 to 12 months and 1,374 genes were correlated to 12 month cortical surface area. Of those 1,227 genes were correlated to both phenotypes (**Figures 5a,b**). Those brain-size-correlated genes in sc_22 at day 14 were enriched in genes involving RHO GTPase Effectors and Cell Cycle pathways (**Figure 5c, Table S7**) including *BUB1*, *KLF14*, and *TUBA1A* which has previously been implicated in outer RG mitotic and migration behaviors^52^ (**Figure 5d**). This observation prompted a further investigation to determine the developmental stage of sc_22 at day 14 (**Figure 1d**). We employed pseudotime analysis to investigate the progression of these cells (**Figures 5e, S7**). The analysis revealed that although sc_22 clustered within a neuronal class when considering cells from both time points, day 14 sc_22 is a cell type at the convergence of fate decision between progenitor and neuron, supporting the radial unit hypothesis where fate decisions toward progenitor cells expand the progenitor pool and lead to increased neuronal production and cortical surface area^21^. We next explored the potential relevance of brain-size-related genes to other characterized phenotypes. We performed an enrichment analysis on multiple gene sets including those known to impact brain size: (primary) microcephaly and macrocephaly. We found that genes associated with brain size, especially those positively correlated, were enriched in microcephaly associated genes, providing additional support for the function of these genes in determining cortical size (**Figure 5d,f**)^53–55^. Furthermore, genes associated with risk for ASD, developmental delay (DD), and neurodevelopmental disorders (NDD)^56^, were also enriched in genes correlated with brain size suggesting a convergence between the processes of early brain growth and risk for neurodevelopmental disorders (**Figure 5f**). Overall, we find that gene expression variability within a cell type at the crux of a crucial fate decision between progenitors and neurons was strongly associated with the size of the infant brain, implicating early neural progenitor fate decisions in determining cortical surface area.

**Fig. 5.**
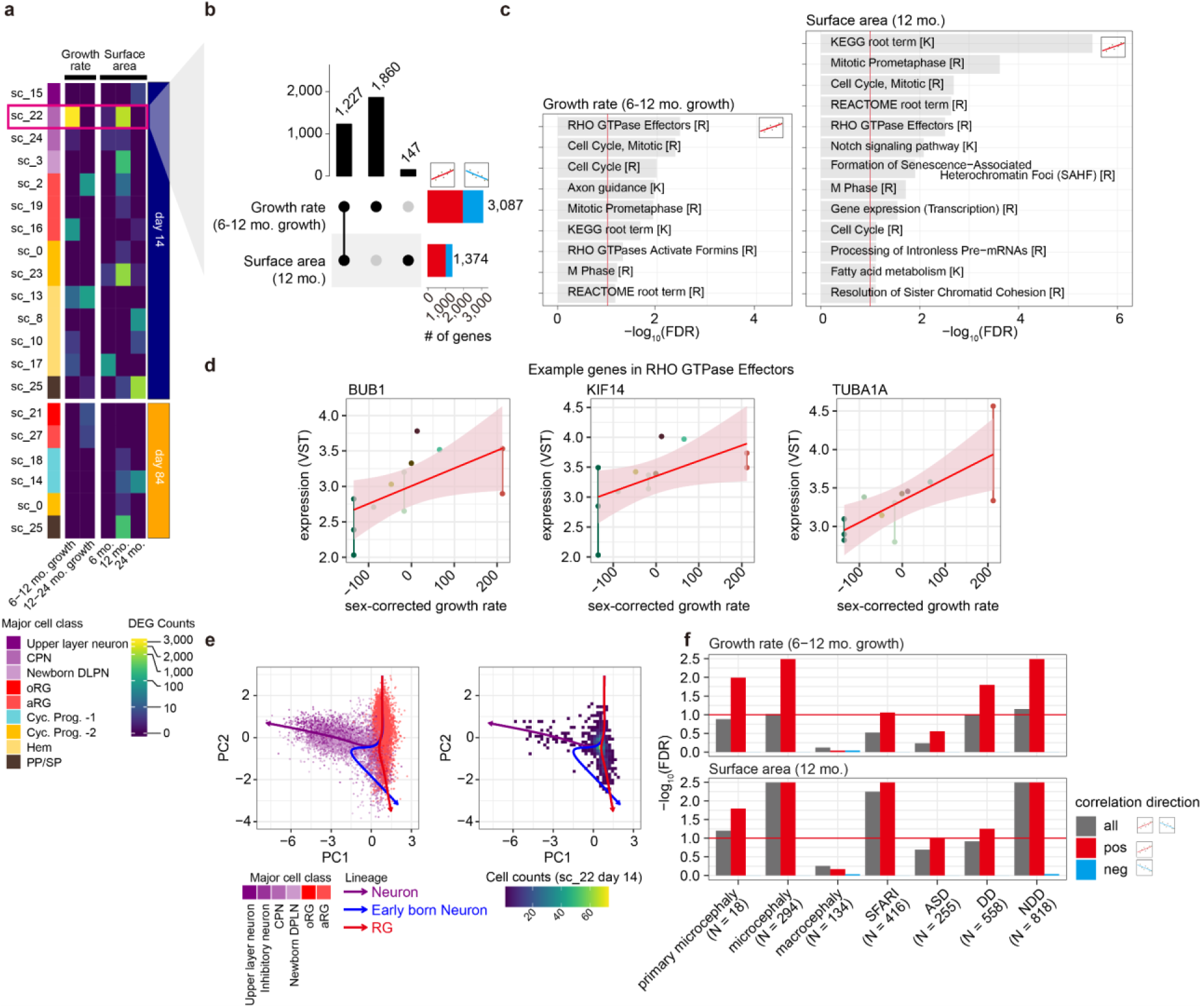
Gene expression levels are associated with surface area and growth rate. **a**, Heatmap showing the distribution of gene counts associated with brain growth rate (left) and surface area (right) across subclusters and time points. Color intensity indicates the number of associated genes. In the subsequent panels, we focus on genes associated with sc_22 at day 14 for growth rate (6-12) and surface area at 12 months. **b**, UpsetR plot summarizing gene overlaps, **c**, Pathway enrichment analysis of associated genes, with key pathways such as “*RHO GTPase effectors”*, “*Cell Cycle”*. **d**, Scatter plots demonstrating correlations between sex-corrected growth rate and gene expression (three examples from RHO GTPase effectors). **e**, PCA of gene expression from subset major cell types (Upper layer neuron (ULN), Inhibitory neuron (IN), cortical projection neuron (CPN), newborn deep layer projection neuron (DPLN), outer radial glia (oRG), apical radial glia (aRG)) colored by cell class (left). Extracted cells from sc_22 on the same dimension in the left panel. Color intensity indicates cell count density (right). Three lineages (ULN: purple, newborn neuron: blue, RG: red) were annotated using inferred pseudotime. **f**, Genes positively associated with brain growth or surface area were enriched in microcephaly and ASD.

## Discussion

Using iPSC-derived hCOs from deeply phenotyped participants with longitudinal MRIs during infancy, we identified *in vitro* phenotypes that predicted infant brain growth. Variability in the expression of cell cycle and RHO GTPAse genes at early time points in development when progenitors are undergoing fate decisions between self-renewal and neurogenesis, as well as the proportion of cortical hem cells, were associated with cortical surface area growth rates during infancy. hCO size during early differentiation was also nominally associated with later cortical surface area in the same individual. Together these data suggest that iPSC-derived hCOs can model inter-individual variation in cortical surface area in the absence of large effect size variants.^12,13,57^

Our data set utilized multiple clones and technical replicates from the same participant, allowing us to capture variability due to reprogramming. We found that genetic background influenced morphology and cell type composition, as shown by higher similarity from the same participant than from other participants. While the use of multiple clones is recommended to remove poorly differentiating clones, this guideline is infrequently practiced due to the considerable expense^58^. Our sample size of 18 participants exceeds previous studies that compared iPSC-derived cortical progenitor cells to directly measured participant brain size^59^ and is comparable to those that nominally bin brain size into macrocephaly and typically developing groups^17^. Relatively large sample sizes of iPSC-based studies are likely necessary for mapping *in vivo* to *in vitro* relationships in the absence of large effect size variants^60^. Finally, utilizing the uniquely extensive phenotyping of this IBIS data set, we found that longitudinal cortical surface area growth rates, rather than cross-sectional snapshots of brain size, were most often associated with *in vitro* cellular and molecular phenotypes, likely because infant brain growth is highly dynamic and growth rates normalize some variability in cross-sectional measurements.

Most hCO phenotypes associated with cortical structure were detected early in differentiation (day 14). We propose two non-mutually exclusive interpretations of this finding: 1) decreased variability due to less time in culture increased the likelihood of identifying *in vitro* correlates to brain size and 2) early progenitor pool expansion has a strong effect on increasing the neurogenic output and cortical surface area, potentially through generating larger populations of IPs or oRGs (**Figure S6e**). Though hCO models have increased in complexity and are routinely grown for extended periods, our results emphasize the importance of collecting early phenotypes that may be more relevant to participant cortical growth.

During fetal development the cortical hem is a major organization center that secretes WNT and BMP to help shape the developing cortex^61^. WNT signaling is known to influence progenitor proliferation, and genetic variation within WNT responsive regulatory elements influences cortical surface area^49,50^. We found that an increase in the proportion of cortical hem cells is associated with cortical surface area growth rate from 6-12 months. In addition, we also identified a positive association of WNT signaling genes in a cortical hem subcluster (sc_8) with 24 month cortical surface area. These findings are similar to previous research that identified a correlation between iPSC-derived neural progenitor cell proliferation rates and participants’ total brain volume. The same study reported that WNT modulation of iPSC-derived neural progenitor cells could attenuate proliferation rates in ASD participant derived neural progenitor cells to typically developing participant levels^59^. hCOs are considered to have endogenous WNT signaling that influences their differentiation outcomes, usually by changing the proportions of cortical neurons^17,62^. These findings support the hypothesis that endogenous WNT signaling impacts cortical size.

We also identified a correlation between 6-12 month cortical surface area growth rates and RHO GTPases and cell cycle genes in sc_22, a subcluster of CPN. Though RHO GTPases have known roles in dendrite, axon and synapse development^63^, they also play a key role in neural progenitor morphology, proliferation, and cell fate decisions^52,64,65^. Pseudotime trajectory suggests this subcluster represents early progenitors undergoing cell fate decisions between self-renewal and neurogenesis. There are multiple lines of evidence suggesting that progenitor cell fate decisions alter cortical surface area: cortical surface area heritability is enriched in regulatory elements of neural progenitors^66^, cell fate decisions within the first several weeks of organoid differentiation have been shown to correlate with brain size differences across species^67^, and previous studies of hCOs identified a positive correlation between progenitor cell cycle and total brain volume of participants from which iPSCs were derived^59^. The greater number of associations detected with earlier (6-12 month) cortical surface area growth rates may be due to their stronger genetic control at earlier ages^68^. These data provide additional support that progenitor cell fate decisions in early development are a critical process for determining cortical size.

### Limitations of the Study

We acknowledge that hCOs have missing cell types including astrocytes, oligodendrocytes, microglia, and vasculature, and are modeling early to mid-fetal developmental processes that make them incomplete models of cortical development. Nevertheless, we did identify relationships specific to progenitor and cortical hem behavior, indicating that hCOs predicted cortical size of the individual from which they were derived. Though hCO model systems are imperfect, we demonstrate that these cell types do show detectable fidelity. Our sample size was not adequately powered to perform group-wise comparisons between participants with and without ASD, but the fidelity demonstrated here provides a strong rationale for these questions in future larger studies.

## Methods

### IBIS Participant Recruitment and Blinding

The study, blood draw, and iPSC generation were IRB-approved under a single IRB at UNC that covered collections from multiple sites(IRB number: 17-1871). IBIS participants with at least 2 MRI scans at either 6, 12 or 24 months of age were recruited for iPSC generation during their school age visit at the University of North Carolina at Chapel Hill or Washington University in St. Louis^18,69,70^. Blood draws were conducted after parental informed consent and assent of the child IBIS participant. 7 females and 11 males, all identified as white except for 1 female participant identifying as multiracial. 4 participants received an ASD diagnosis at 2 years of age and had a family history of ASD. Diagnoses were made based on expert clinical judgment using DSM-IV-TR criteria^71^ and all available clinical information, including direct examination using the Autism Diagnostic Observation Scale^72^, parent report during Autism Diagnostic Interview-Revised^73^, and other behavioral measures. 9 participants had a family history of ASD but no diagnosis and 5 participants had no family history or diagnosis of ASD. Blood draws were preserved in sodium citrate (Fisher Scientific 369714) and processed within 48 hours. PBMCs were isolated by Ficoll gradient and cryopreserved at the University of North Carolina’s Human Pluripotent Stem Cell Core. Each participant was assigned a unique ID that is associated with previous IBIS studies, but blinds experimenters to diagnostic status and other criteria during handling of the cell lines.

### MRI Processing

MRI processing was performed as previously described ^18^ and blinded to sex, family history, and participant diagnostic information. Briefly, a CIVET workflow was used to obtain cortical surface area measurements^74,75^ with adaptations to infant ages for automated anatomical labelling^76,77^. Multi-model, within-subject co-registration of 12 mo. cortical surface area was used to extract data from 6 mo. cortical surface area. All surface area measurements were quantified at the mid-cortical surface, and absolute growth rates were used, relative to normative body growth rather than chronological age^18^.

### Reprogramming and iPSC Validation

Each participant had 3 iPSC clones reprogrammed from their PBMCs. Except for the first participant, reprogramming was performed in batches of 4 participants, with at least one participant from every diagnostic group represented. Erythroblasts were reprogrammed using a non-integrating Sendai virus based method (CytoTune-iPS 2.0)^27^. Following reprogramming, individual colonies were selected and passaged onto mouse embryonic fibroblasts and then transitioned into feeder free conditions on hESC qualified matrigel (Corning 354277) and mTeSR+ (STEMCELL Technologies 05825). The Human Pluripotent Stem Cell Core mainted a database of experimental details that could impact the clones (See hCO Differentiation). Pluripotency of each iPSC clone was confirmed using immunocytochemistry of pluripotency markers and tri-lineage differentiation followed by qPCR assessment of marker gene via Taqman ScoreCard. Each clone also underwent PCR to test for mycoplasma contamination before banking the iPSCs at low passage number (passages 4-6), and were found to be negative. All iPSC lines were transitioned to and maintained in mTeSR+ on hESC-qualified matrigel coated plates. Versene (Thermo Fisher 15040066) was used to passage cells. Mycoplasma testing using PCR detection was performed before each experiment and only mycoplasma negative iPSC lines were used for subsequent differentiations.

To confirm the iPSC clone’s participant identity, the participant’s genotype from a previous IBIS project (Illumina HumanOmni5-Quad) or from PBMCs isolated for this study (Illumina Infinium GDA-8 v1.0 kit #20031810) were compared to genotype information for each clone (Illumina Infinium GDA-8 v1.0 kit #20031810). Potential sample swaps were detected by identity-by-descent (IBD) analysis using plink (v1.90)^78^ on both iPSC gDNA and from transcriptomes from scRNAseq. Reassignment of participant’s ID to iPSC clone was verified using DNA isolation from both PBMCs and iPSCs followed by comparing lanes of agarose gels with amplified short tandem repeats^79^. No aneuploidy was detected by visualizing Log-R ratios and B-Allele frequency in Genome Studio 2.0 (**Figure S1**). When sample swaps were detected in scRNAseq transcriptomics, imaging data from samples with mislabeled identity were removed from further analysis. Sample swaps detected at day 14 or in one differentiation of an iPSC clone was sufficient evidence to exclude all data from that clone.

### hCO differentiation

The organoid differentiation protocol was adapted from to feeder-free conditions as previously described^4,30^. After at least 2 passages of the iPSC line, organoids were aggregated into wells of 96 well v-bottomed ultra-low attachment plate (S-Bio MS-9096VZ) in recovery media (6 μM Y27632 (Fisher Scientific ACS-3030) in mTeSR+). Nine thousand singularized iPSCs (Accutase Thermo Fisher A1110501) were used per well in 200 μL recovery media. At day 0, 100 μL was replaced with Forebrain First Media (DMEM/F12 Thermo Fisher 11320033, 20% Knock-Out Replacement Serum Life Technologies 17502048, Glutamax Life Technologies 35050-061, MEM-Non Essential Amino Acids Gibco 11140050, Pen-Strep Gibco 15140122, Beta-Mercaptoethanol Thermo Fisher Scientific 21985023) with double the concentration of SMAD inhibitors (8uM Dorsomorphin Sigma-Aldrich P5499, 8uM of A-83 STEMCELL Technologies 72022). From day 1 to day 4, 150 μL of media was replaced with Forebrain First Media (4uM Dorsomorphin, 4uM A-83). Day 6 and day 7, wells were fed with 100 μL Forebrain Second Media (DMEM/F12 Thermo Fisher 11320033, N2 Life Technologies 17502048, Glutamax Life Technologies 35050-061, MEM-Non Essential Amino Acids Gibco 11140050, Pen-Strep Gibco 15140122, Beta-Mercaptoethanol Thermo Fisher Scientific 21985023) with a SMAD inhibitor (SB-431542 Fisher Scientific 16-141) and WNT activator (CHIR-99021 Fisher Scientific S12632). At day 7, organoids were embedded in a 60% Growth Factor Reduced Matrigel (Corning 356230) and Forebrain Second mixture in a well of an ultra low attachment six well plates (Sigma-Aldrich CLS3471), and then cultured with 3 mL of Forebrain Second media. From day 8 to day 13 cultures were fed every other day with Forebrain Second media.

On day 14, all organoids were removed from the embedding and transferred into non-tissue culture treated 12 well plates, ∼10 per well. Each well was fed with 4 mL of Forebrain Third Media (DMEM/F12 Thermo Fisher 11320033, N2 Life Technologies 17502048, Glutamax Life Technologies 35050-061, MEM-Non Essential Amino Acids Gibco 11140050, Pen-Strep Gibco 15140122, Beta-Mercaptoethanol Thermo Fisher Scientific 21985023, B-27 Life Technologies 17504044, 2.5ug/mL Insulin Sigma-Aldrich I9278-5ML). Miniaturized spinning bioreactor (SpinΩ bioreactor) with the DC motor set to ∼100 rotations per minute of the internal paddles (7.5 V) were used as lids. Organoids were fed with Forebrain Third Media every 2 to 3 days. Day 35 and after, F3 media was supplemented 1:100 with growth factor reduced matrigel. Organoids were separated out into new wells if the media color changed rapidly or if organoids were developing cysts. From day 70 to day 84, organoids were fed with Forebrain Fourth Media (Neurobasal Life Technologies 21103-049, Ascorbic Acid Sigma-Aldrich A4544, Glutamax Life Technologies 35050-061, MEM-Non Essential Amino Acids Gibco 11140050, Pen-Strep Gibco 15140122, B-27 Life Technologies 17504044, cyclic AMP 0.5mM Sigma-Aldrich A6885, 20ng/mL BDNF Life Technologies PHC7074CS1, 20ng/mL GDNF R&D Systems 212-GD-010).

All clones passing quality control were randomized to 5 initial rounds of differentiation so that each round contains participants from different diagnostic groups and from different reprogramming batches. These rounds were thawed at the same time and usually seeded together. Technical variables associated with reprogramming and the differentiation were recorded in a database and included the following: Site, Collection_Date, Mail_Hand, Mailed_Date, Date_Received, Received_Initial, Blood_Processing_Date, Blood_Processing_Initial, Isolation_Experimenter, Blood_mixing_with_Ficoll, Number_of_Viable_Cells_Millions, Number_of_frozen_PBMCs_vials, Number_of_cellsvial_Millions, ReprogrammingBatch, PBMCID, Thaw_date_for_PBMC_vial, QBSF_Lot_Number, L-3_Lot_Number, EPO_Lot_Number, IGF-1_Lot_Number, Dexamethasone_Lot_Number, MOI_added, CytoTune_Lot_Number, Person_Reprogramming, Day_colonies_appear, DMEMF12, KSR, L-glutamine, MEM_NEAA, FGF_Lot_Number, Number_of_iPSC_colonies_picked_for_P0, P0_Selection_Date, Lift Minutes, Seeding_Minutes, SeedingViability, F1_Start, Plate Location, Count_D0, PlateID, Count_D7, Embedding_Minutes, A83, Accutase_Lot, Round, Freeze_date, Thaw_Date, Split_Ratio_Set_1, Split_Ratio_Set_2, Day0_Date, Well_Pair_for_seeding, Additional_Well_Pair_for_seeding, F3 Start Date,NumberofWellsDay14, CountD14, CountD35, CountD56, CountD70, CountD84, LotNEAA, BiorectorFailure, PassagetoDiff.

An additional 4 rounds of differentiation were completed to include an additional participant and perform a second differentiation of 27 of cell lines where the first differentiation failed due to hCO death before day 14, no organoids with visible budding by day 14 or < 20% PAX6 expression in intact immunolableing.

### qPCR for iPSC cell state

300,000 iPSCs were frozen in Trizol stored at -80℃ from the same iPSCs as those seeded. in 6 to 8 iPSC lines had total RNA extracted using the RNeasy columns (Qiagen 74106). cDNA was generated using the iScript cDNA synthesis kit (Bio Rad 1708891). qPCR reactions used SsoAdvanced Universal SYBR Green Supermix (Bio-Rad) and a QuantStudio 5 Real-Time PCR System (Thermo Fisher). We considered PGP1 the control cell line for calculating ΔCT. Gene Expression was normalized to *EIF4A2* expression. Primers: *EIF4A2* (Forward: TGGTGTCATCGAGAGCAACTG, Reverse: GGCTTCTCAAAACCGTAAGCA), *NANOG* (Forward: TTTGTGGGCCTGAAGAAAACT, Reverse: AGGGCTGTCCTGAATAAGCAG), *OCT4* (Forward: GGAGAAGCTGGAGCAAAAC, Reverse: ACCTTCCCAAATAGAACCCC), *OTX2* (Forward: AGAGGACGACGTTCACTCG, Reverse: TCGGGCAAGTTGATTTTCAGT), *ZIC2* (Forward: GATGTGCGACAAGTCCTACAC, Reverse: TGGACGACTCATAGCCGGA), *DUSP6* (Forward: TTCCCTGAGGCCATTTCTTT, Reverse: AGTGACTGAGCGGCTAATG), *KLF4* (Forward: GCTGCCGAGGACCTTCTG, Reverse: GCGAACGTGGAGAAAGATGG), *TFAP2C* (Forward: CTGTTGCTGCACGATCAGACA, Reverse: CTCAGTGGGGTTCATTACGGC), *TFCP2L1* (Forward: ATACCAGCCGTCCTATGAAACC, Reverse: ACTGCGAGAACCTGTTGCG), *SOX2* (Forward: GCTACAGCATGATGCAGGACCA, Reverse: ATCGCCCCACTTGATTTTGG), *SOX17* (Forward: CTCTGCCTCCTCCACGAA, Reverse: CAGAATCCAGACCTGCACAA), *SNAI2* (Forward: CGAACTGGACACACATACAGTG, Reverse: CTGAGGATCTCTGGTTGTGGT)

### Phase Contrast Imaging and Analysis

Phase contrast images of the organoids at day 0, 7, 14, 35, 56, 70 and 84 were captured on an EVOS XL Core microscope with a 10x or 4x objective. Automatic cross-sectional area measurements were acquired using Cell Profiler customized pipelines for each time point. Manual tracing of cross-sectional area was performed blinded to any participant information where necessary. Three independent rankers were trained on previously used organoid morphology rank scale until 85% inter-rater agreement on a test set.^4^ Ranking and area measurements were performed blinded to the participant’s diagnostic or cortical surface area data. hCOs without visible buds were excluded from correlations to MRI measurements. The largest and smallest 2.5% of area measurements were excluded from area measurements as outliers. Additional outliers for each participant were identified if the sample was outside the inter quartile range for that measurement. Correlations to MRI measurements were performed using average area measurements for each participant and controlling for sex in a linear regression model. Age of MRI was controlled for in the cross sectional MRI measurements. FDR adjustment using Benjamini-Hochberg (BH) procedure was applied to correct for each area measurement to all MRI measurements.

### Immunocytochemistry

For 2-D immunolabeling, organoids were sectioned into 12 microns slices using a cryostat and mounted onto slides and immunolabeled as previously described^4^, coverslipped in Fluoromount-G mounting media with DAPI (Thermo Fisher 00-4959-52) and images on an Olympus FV3000RS with a 20x objective. The following primary antibodies were used: CTIP2(1:200, AB_18465), KI67 (1:200, AB_10853185), TTR (1:200, AB_2804646), EOMES (1:200, AB_2572882), HOPX (1:500, AB_10603770), PH3 (1:200, AB_10984484).

### Tissue Clearing and Imaging

Day 14 organoids were tissue cleared using an adapted iDISCO+ protocol as previously described^4,80^. Briefly, individual hCOs were permeabilized, incubated in a blocking buffer and then transferred to individual wells of a 96 well plate. Organoids were incubated in primary antibodies for 18 hours, while on an orbital shaker at 90 r.p.m. PAX6 (1:100 AB_2792516), NCAD (1:200, AB_398236). After several washes, hCOs were then incubated with TO-PRO-3 (Thermo Fisher, T3605, 1:300) and secondary antibodies were added for at least 18 hours at 37℃ with agitation. Samples were then washed and embedded in 1% agarose blocks between 2-5mm thick, where relative location was used to keep track of sample identity. Agarose blocks of organoids were dehydrated and placed in dibenzyl ether solution (RI = 1.56, Sigma-Aldrich, 108014) for imaging within 3 days of completion of the staining protocol. Day 14 organoids (<1mm) were imaged on a Olympus FV3000RS, using an Olympus LUCPlanFL-N 20x long working distance objective and Galvano scanning. 3-5 organoids were imaged per clone, with up to 3 clones per agarose block.

ImarisFileConverter (v9.9.0 or 9.7.0) and IMARISStitcher (9.10.0 or 9.7.0) were used to preprocess the Olympus images. Surfaces were created with manual thresholding and classification while blind to participant information for ToPRO, NCAD, and PAX6. Correlations to MRI measurements were performed using average measurements for each participant and controlling for sex in a linear regression model. Age of MRI was controlled for in the cross sectional MRI measurements. FDR was applied to adjust for multiple testing across comparison of each volumetric measurement to all MRI measurements.

### Single Cell Dissociation and Library Preparation

Organoids were dissociated using papain (Worthington Biochemical Corporation LK003150)^6^. At day 84, a phase contrast image was taken of each organoid harvested for scRNAseq. These organoids were selected to have smooth edges and no visible cysts. At day 14, organoids were selected randomly for dissociation. All scRNAseq samples had no more than 30% dead cells in Trypan blue counting. Single cells were permeabilized and fixed using ParseBiosciences Cell Fixation Kits (SB1001). Barcoding and library preparation with Whole Transcriptome Kits (WTK) (EC-W01030, v.1.3.0), a split-pool approach uses random hexamer and Oligo-dT barcodes to label individual cells. Each kit except for the first contained samples from multiple rounds of reprogramming and differentiation with a mix of IBIS participant diagnostic status.

### scRNA-seq data processing

Libraries were sequenced with i7 Illumina primers on a Novaseq 6000 S4 flowcell with 10% Phi-X, using 100 or 150 base pair paired-end reads. Random hexamer barcodes in the sequencing reads were converted to paired Oligo-dT barcodes to create the final reads derived from the same cell using *splitp* (v1.0) (https://github.com/COMBINE-lab/splitp). Reference containing spliced transcripts and intronic sequencing was created using *make_splici_txome*() of roe package (https://github.com/COMBINE-lab/roe). After excluding low quality reads (phread quality score < 20), reads were aligned to the reference using salmon alevin^81^ (v.0.6). Gene expression matrices were generated by alevin-fry (v0.5.1) ^82^ and imported into R using *loadFry*() of fishpond (v2.0.1)^82^ with snRNA mode, which covers the spliced, unspliced and ambiguous mRNA count of genes. Data was initially processed using the Seurat R package (version v4.0.3)^83^ standard pipeline.

### Quality control on scRNA-seq data

Cells were filtered out if (1) gene < 1,000 per cells or (2) RNA expression counts < 1,500 per cells or (3) a mitochondrial gene expression percentage >=10% of total RNA expression. Data was split to a unique sample defined as a combination of participant’s ID, batch, Whole Transcriptome Kit (WTK1-5), and organoid differentiation day (14 or 84). We estimated doublets within sample using doubletFinder (v2.0.3) (https://github.com/chris-mcginnis-ucsf/DoubletFinder)^33^ with an expected doublet rate (3.4% or 4.25% for WTK1-3 and WTK4 and 5, respectively) and pN = 0.25. Optimal pK was set to maximum of the bimodality coefficient distribution. At this stage, we obtained 210,016 cells in total (median = 995 per sample, s.d. = 2,323).

Because of memory issues due to the large dataset, the following analyses were performed by Seurat V5^35^. *SCTransform* (v2) was applied separately to each sample for transformation and normalization to correct batch effect prior to integration by *IntegrateLayers*(method=RPCAIntegration,k.anchor=20). Cell type clusters were defined by *FindNeighbors*(), followed by *FindClusters*(). We obtained 35 cell clusters by choosing a resolution of 1.0.

To identify and exclude stressed cells, we applied granular functional filtering (Gruffi)^34^. The detailed instructions are provided in their GitHub repository (https://github.com/jn-goe/gruffi). Briefly, we partitioned cells into groups using a clustering function implemented in Gruffi, and calculated Gene ontology scores of (*Glycolytic process*, *ER stress*, *Gliogenesis*) per cell, and averaged them per granule. We defined cells as stressed cells if the Glycolytic process or ER stress cells score were high but Gliogenesis scores were low using a threshold of 90th percentile. We removed stressed cells and retained only IBIS participant derived iPSC hCOs for downstream analyses.

### Cell Type Annotations

Cell type clusters were grouped by comparing marker genes in publicly available primary fetal tissue data^36^. First, we ran *FindAllMarkers*() to identify marker genes in our dataset (FDR < 0.05; avg_log2FC > 0.25, pct.1 > 0.25 | pct.2 > 0.25), then performed Fisher’s Exact test using gene markers from our clusters and their marker genes on all cluster combinations. The odds ratio and significance of enrichment were estimated using a two-sided Fisher’s exact test, followed by multiple test correction (significant threshold at FDR < 0.05). Next, we performed hierarchical clustering using the *p*-values for all pairs obtained in the previous step. Finally, the optimal number of clusters (12 = cell class; K = 12) was determined based on the Scree plot. Each of the 12 cell classes were then manually annotated based on marker gene expression and comparison to *in vivo* tissue.

### Cell Type Proportions

Cell type proportions for cell classes and subclusters were calculated for each differentiation at day 14 and day 84 following the removal of stressed cells and samples identified as outliers. Proportions were calculated using Propeller’s default logit transformations^84^. Unknown cell types were removed for correlating cell type proportions to participants’ MRI measurements. Correlations to MRI measurements were performed using average measurements for each participant and controlling for sex in a linear regression model^85^. Age of MRI was controlled for in the cross sectional MRI measurements. FDR correction was applied for each cell class proportion at day 14 or day 84 correlation across all MRI measurements, and was implemented separately for subclusters and cell classes.

### Correlation to technical variables

Technical variables associated with organoid outcomes were identified using linear models. Each differentiation is considered one sample. Area measurements and cell type proportion data had undergone quality control as described above. Unknown cell types were included for this analysis. FDR was calculated for cell type proportion correlates to technical factors by correcting over all technical variables across all subclusters and cell classes (92 classes and subclusters, 78 technical variables). FDR was calculated for each differentiation’s average cross-sectional area measurements, hCO morphology, and intact immunolabeling measurements together (14 measurements, 78 technical variables).

### Cross assay correlations

Pearson’s correlation coefficient was used to calculate cell class proportion correlates to each differentiation’s average cross-sectional area measurements (N = 54), hCO morphology (N = 54), and intact immunolabeling measurements (N = 19). All hCOs from a given clone within a differentiation batch were averaged and considered as one sample. Area measurements and cell type proportion data had undergone quality control as described above.

### Pseudo-bulk differential gene expression analysis

Per each subcluster-clone-day, we aggregated gene counts to generate pseudo-bulk expression matrices by *aggregateData*() implemented in muscat (v1.13.1)^86^. The expression values were then normalized by Variance stabilizing transformation (VST) (DESeq2 v1.38.3)^87^. Because one participant can have multiple clones, we estimated the within-subject correlation across participants using *duplicateCorrelation*() within limma (v3.54.2)^88^. By incorporating the correlation structure, VST counts were fit to a linear model with sex (and age at visits) as covariates while adjusting for intra-individual similarities, followed by empirical Bayes moderation. For each MRI measurement and pseudo-bulk dataset, we excluded clones from the analyses if (1) MRI measurement was not available for the corresponding participant, or (2) if pseudo-bulk was calculated from < 20 cells or genes were detected in less than 10% of individuals in the subcluster-day group. subcluster-day groups representing less than 7 participants were also excluded. To avoid potential outlier influence on the results, we calculated Cook’s D and removed the gene-participant combination if Cook’s D > 4/N where N is the total number of clones included in the analysis, and then repeated limma analysis. FDR was used for multiple test corrections for each subcluster-day and phenotype.

### Pathway enrichment analysis

KEGG or REACTOME Pathways^89,90^ enriched in subclusters correlated to any MRI measurements were identified by g:profiler(v0.2.3)^91^ on genes found by *FindMarkers*() of Seurat. In this analysis, we set cells.1 = cells in target subcluster, cells.2 = all cells grouped in different cell classes. Background genes were all genes in the scRNA data. For both marker genes and background genes, we excluded non-protein-coding genes or genes located within MHC region (chr6:28,510,120-33,480,577) from the analysis. FDR< 0.1 was used as the significance threshold.

### Pseudotime analysis

SCT normalized gene counts were extracted from subclusters in the following day 14 cell classes: upper layer neuron, IN, newborn DLPN, CPN, oRG, and aRG. We excluded sc_27, 28, and 33 as those clusters had < 100 cells, resulting in 10 subclusters being used. Genes expressed in a small proportion of cells (<50) were then excluded and fed into PCA analysis followed by slingshot^92^ with default settings.

### Enrichment analysis using genes associated with brain size

Genes associated with MRI measurements (DEG) were selected per subcluster-day and MRI phenotype. Those gene sets were grouped as “Positive”, “Negative”, or “all” depending on the direction of correlation. When the gene set group has at least 50 genes (N_DEG_ >= 50), we performed brain-disorder-related gene set enrichment analysis. To evaluate whether the observed overlap of DEG (O_obs_) and target gene sets is significantly more than expected by chance, we performed a permutation test. Specifically, we selected random sets of genes (N_random_ = N_DEG_) from background genes (= all genes used in DEG analysis in each cluster-day). We repeated 10,000 samplings and generated a null distribution of overlap values (O_rand_1 …._ O_rand_10000_). The *P*-value was estimated as fraction of random overlaps greater than or equal to the observed value. *P*-values were corrected using the BH procedure. We note that we set *P* as 0.0001 for multiple corrections when the fraction of O_rand_ was 0.

Target gene sets and data sources are described as below:

1. Rare variant association study^56^: Autism spectrum disorder (ASD), developmental delay (DD), and all neurodevelopmental disorders (NDDs). (2) SFARI, FDR < 0.1 from the transmission and *de novo* association (TADA) tests were extracted (N = 255, 599, 819, respectively).
2. SFARI Gene (https://gene.sfari.org/): Version: 08-19-2024 Release We chose genes with a gene score < 2 or syndromic. (N = 416)
3. DECIPHER (https://www.deciphergenomics.org/): Microcephaly genes: annotated with HP:0000252 (N = 299) Macrocephaly genes: annotated with HP:0000256 (N = 134)
4. OMIM (https://www.deciphergenomics.org/): “microcephaly” and “primary” were used as queries for gene map retrieval. *Gene Map Search - ‘microcephaly and primary (Search in: Entries with: Genemap; Search in: Title; Retrieve: gene map)’* We ensured that the phenotype column has “microcephaly X, primary” and also excluded provisional cases. (N = 19).

## Acknowledgments

We are sincerely grateful to all the families and children who participated in the Infant Brain Imaging Study that enabled this work. This work was supported by the NIH (3-P50-HD103573-03S1, R01 MH130441, R01 MH121433, R01 MH120125, R01 MH118349), UNC TraCS (550KR262111), and a Research Infrastructure Equipment Allocation Grant from UNC School of Medicine to J.L.S; NIH (R01 HD055741) and the Foundation of Hope (A19 – 1040) to J.P. M.R.G. was supported by NINDS T-32 7431-20 and F31MH124427-01A1. The University of North Carolina High Throughput Sequencing Facility is supported by the University Cancer Research Fund, Comprehensive Cancer Center Core Support grant (P30-CA016086), and UNC Center for Mental Health and Susceptibility grant (P30-ES010126). We thank Pablo Ariel at the Microscopy Service Laboratory and Michelle Itano at the Neuroscience Microscopy Core for assistance with imaging. The Microscopy Services Laboratory, is supported in part by P30 CA016086 Cancer Center Core Support Grant to the UNC Lineberger Comprehensive Cancer Center and the North Carolina Biotech Center Institutional Support Grant 2016-IDG-1016. The University of North Carolina’s Neuroscience Microscopy Core (RRID:SCR_019060), is supported, in part, by funding from the NIH-NINDS Neuroscience Center Support Grant P30 NS045892 and the NIH-NICHD Intellectual and Developmental Disabilities Research Center Support Grant P50 HD103573. The UNC mammalian genotyping core is supported by the Lineberger Comprehensive Cancer Center (LCCC), the Center for Environmental Health and Susceptibility (CEHS), the UNC School of Medicine, School of Public Health, University Cancer Research Fund (UCRF), and the North Carolina Biotechnology Center. UNC’s Bioinformatics and Analytics Research Collaborative Core and the Human Pluripotent Cell Core Facility are supported by UNC School of Medicine. The authors would also like to thank the IDDRC organoid working group (Deborah French, Elisa Waxman, Satoshi Yamashita, Kazue Hashimoto-Torii), Fred Gage, Simon Schafer for assistance with organoid differentiation protocols, Young-sook Kim for assistance with scRNA-seq analyses, Chad Chappell, Heidi McNeilly, Shannon Sweeney, Lisa Flake, Lacy Cheers and M. D. Fallin for participant recruitment and genotyping.

## Author Contributions

M.G., N.M., M.Shen, J.G., J.P., and J.L.S. conceived the project. M.G., A.B. M.Shen, J.G., J.P., and J.L.S. designed the experiments. M.G., N.M., J.S., M.Shen, J.G., J.P., and J.L.S. interpreted the results. M.G., N.M., and J.L.S. wrote the paper with input from all authors. N.M. and M.G. performed statistical and bioinformatics analyses. M.G., A.B., N.P. generated hCOs and imaged hCOs. A.B. and A.S.B. reprogrammed iPSC. M.G. performed scRNAseq library preparation, imaged intact hCOs. N.M., A.B. M.G. and K.S. genotyped iPSCs and checked for aneuploidy. M.G. and L.T.D. performed confocal imaging. M.G., I.C. and T.F. curated and analyzed intact immunolabel images. S.A., M.S., E.D., L.T.D., M.Y., K.S., T.F, K.H., ranked hCOs and performed cross-sectional hCO area measurements. K.E. and S.B. performed qPCR. M.L. and J.S. consulted on statistical and bioinformatic analyses. T.S., J.P., A.Estes., S.D., R.T.S., K.Botteron., N. Marrus, Alan E., S.H., M.Styner, R.C.M., D.L.C.., H.V., K.Bennke, L.Z., H.H., J.G., M.Shen, J.P. collected and curated participant data. M.Shen., J.G., J.P., and J.L.S. supervised the study.

## Data Availability

Further information and requests for resources and reagents should be directed to and will be fulfilled by the lead contact, Jason Stein.

