## Supplemental Figures for "Early cell cycle genes in cortical organoid progenitors predict interindividual variability in infant brain growth trajectories"

### Supplementary Information

**Supplementary Figures** **2**
Supplementary Figure 1| iPSC Generation from IBIS Participants 2
Supplementary Figure 2| scRNAseq QC 5
Supplementary Figure 3| Annotation of Cell Classes 6
Supplementary Figure 4| Technical variables associated with cell type proportions 7
Supplementary Figure 5| Reproducibility of cell type proportions and exclusion of poor differentiations 9
Supplementary Figure 6| Intact organoid imaging and technical correlates 10
Supplementary Figure 7| Top variable genes contributing pseudotime trajectory 12
**Supplementary Tables** **13**
Supplementary Table 1| Participant included in each experiment 13
Supplementary Table 2| Marker genes for scRNAseq subclusters and cell classes 13
Supplementary Table 3| Technical correlates to cell classes and subclusters 13
Supplementary Table 4| Cell type and subcluster proportions to MRI measurements 13
Supplementary Table 5| Technical correlates to hCO area and immunolabeling 13
Supplementary Table 6| Genes correlated to MRI measurements 13
Supplementary Table 7| Enriched pathway in DEG 13

### Supplementary Figures

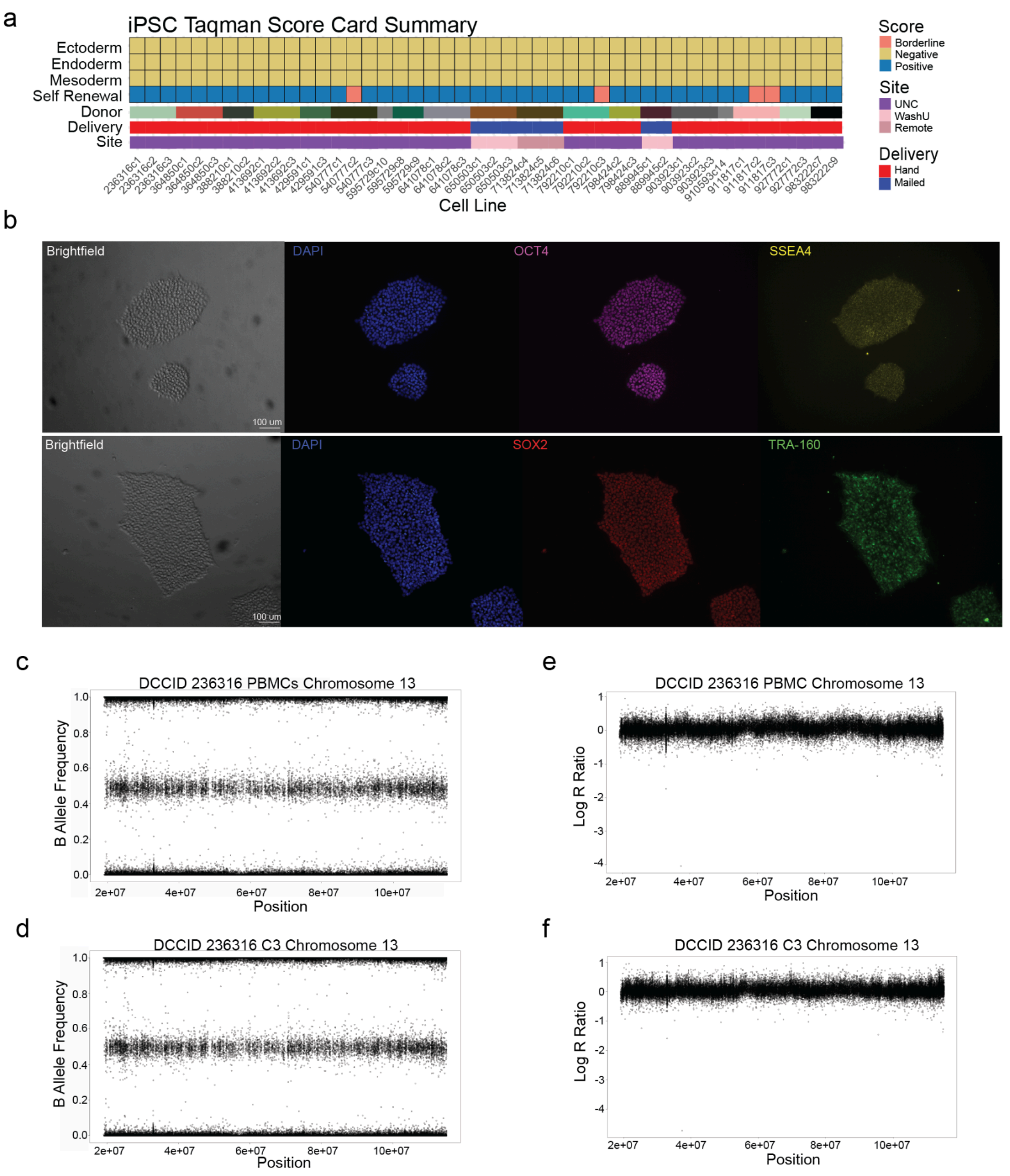


#### Supplementary Figure 1| iPSC Generation from IBIS Participants

**a,** IBIS iPSC Reprogramming. Summarized qPCR Taqman scorecard results for IBIS-iPSC clones in pluripotency promoting media. **b,** Immunofluorescence for pluripotency markers in a representative clone. Example B-allele frequencies (**c**,**d**) and LogR ratio (**e**,**f**) from Illumina global SNP array for one participant's PBMCs prior to reprogramming (**c**,**e**) and one reprogrammed iPSC clone (**d**,**f**), demonstrating no reprogramming induced aneuploidy.

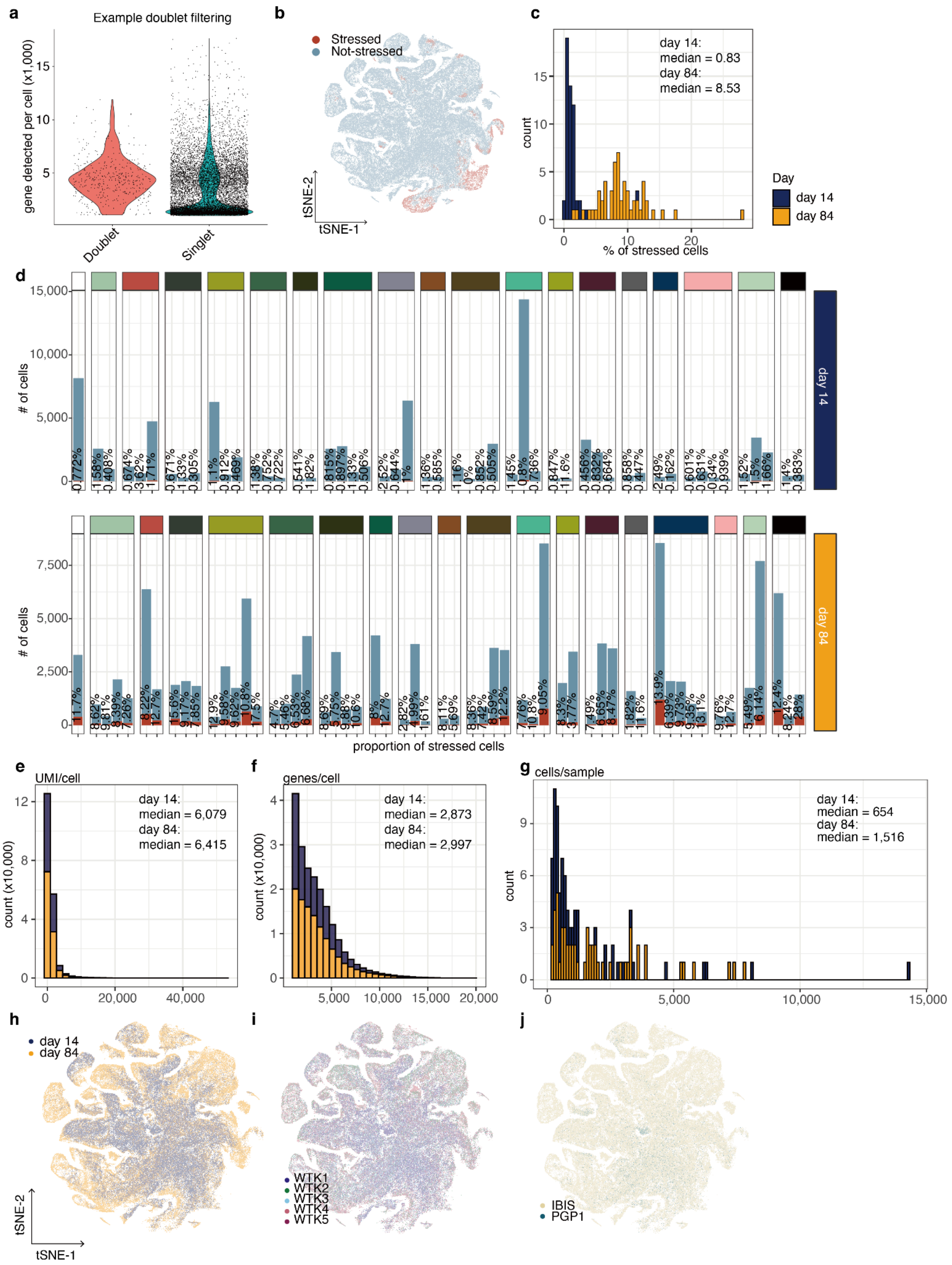
a
Example doublet filtering
gene detected per cell (x1,000)
Doublet
Singlet
b
Stressed
Not-stressed
tSNE-2
tSNE-1
c
day 14:
median = 0.83
day 84:
median = 8.53
count
% of stressed cells
Day
day 14
day 84
d
### of cells
day 14
day 84
proportion of stressed cells
e
UMI/cell
day 14:
median = 6,079
day 84:
median = 6,415
count (x10,000)
f
genes/cell
day 14:
median = 2,873
day 84:
median = 2,997
count (x10,000)
g
cells/sample
day 14:
median = 654
day 84:
median = 1,516
count
h
day 14
day 84
tSNE-2
tSNE-1
i
WTK1
WTK2
WTK3
WTK4
WTK5
j
IBIS
PGP1

### Supplementary Figure 2| scRNAseq QC

**a**, Doublet detection was completed within samples (clone:sequence batch:day triplets). Gene counts detected per cell were higher in doublets, as expected. **b**, Stressed cells on tSNE dimensionality reduction clustering plot, which are mostly in Unknown cell types (55.8% unknown and 24.8% hem). **c,** Histogram showing the stressed cell count at day 14 and day 84. **d**, Stressed cell counts (red) per participant at day 14 (top) and day 84 (bottom) with percentage of stressed cells. Stressed cells were filtered out prior to final analyses. Histograms showing the distribution of **e,** UMI per cell ,**f,** genes detected per cell, and **g,** cells per sample at day 14 and day 84. Cell annotation by **h,** Day, **I,** sequence batch **j,** and either all IBIS samples or a previously published cell line (PGP1) on tSNE plots.

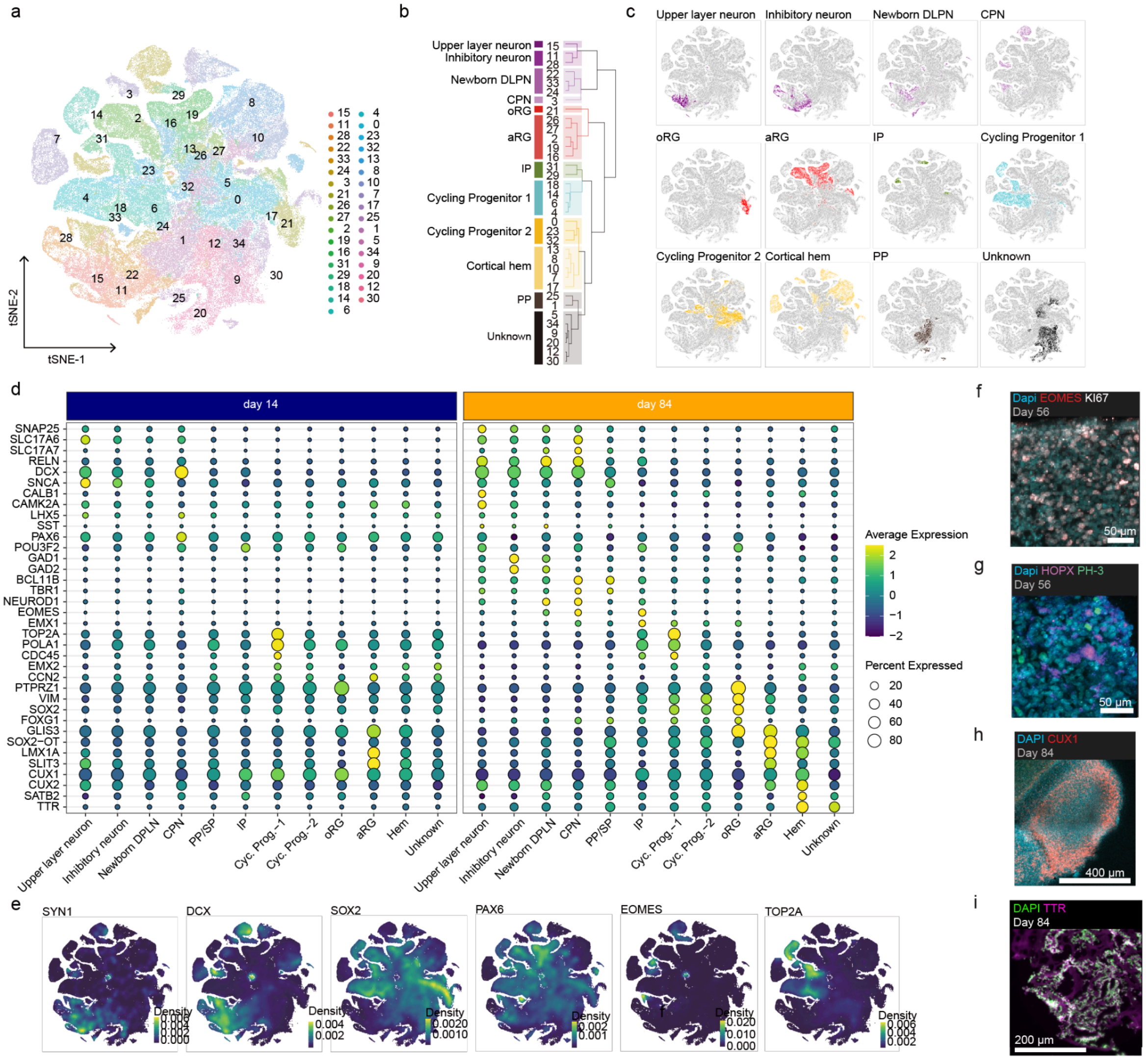


### Supplementary Figure 3| Annotation of Cell Classes

**a**, tSNE plot showing 35 subclusters. **b-d**, Subclusters were then merged to 8 cell classes using hierarchical clustering of transcriptomic profiles, and annotated based on expression of marker genes and comparison with primary fetal tissue. **b,** Dendrogram representing subcluster and cell class. **c,** tSNE plot showing 12 cell classes including those unknown. **d,** Expression of marker genes within cell classes. **e**, density plots of known marker genes. **f**, Day 56 hCO with Ki67+ EOMES+ proliferating intermediate progenitors. **g,** Day 56 hCO with HOPX+ oRG and proliferating PH3+ cells. **h,** Day 84 hCO with CUX1+ upper-layer neurons in a neuroepithelial bud. **I**, TTR+ cells in a day 84 hCO.

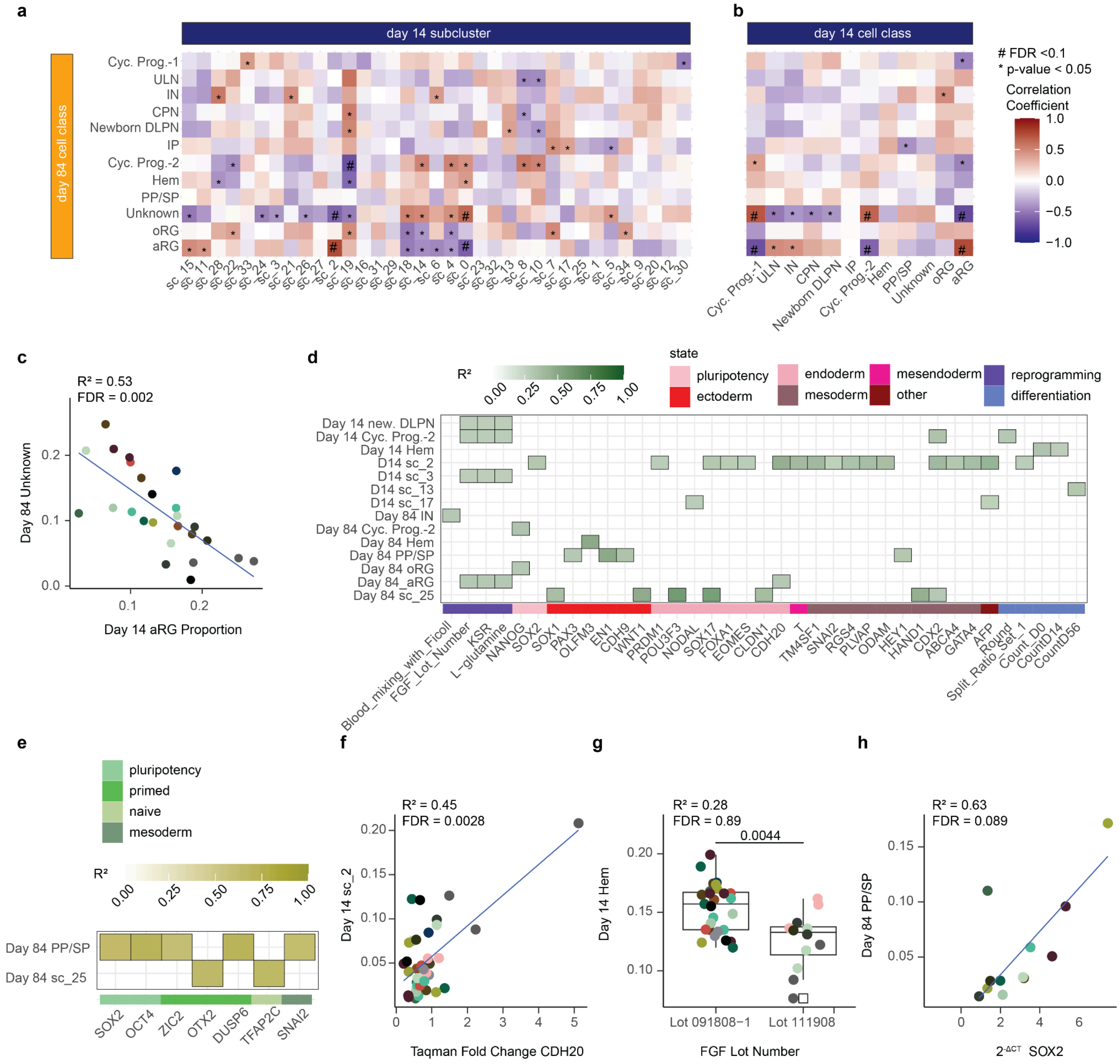

### Supplementary Figure 4| Technical variables associated with cell type proportions

**a,** Pearson's correlations between day 84 cell type proportions for day 14 subclusters and **b,** day 14 cell classes. **c,** Proportions of aRG at day 14 are negatively associated with unknown cells at day 84. **d,** Technical correlates to cell class and subcluster proportions at day 14 and day 84. Technical correlates are shown that survive FDR < 0.1, when correcting over all technical variables across all subclusters and cell classes. Results are shown for all cell classes and subclusters which are shown in the manuscript (N = 92 classes and subclusters, 78 technical variables), day 14 cell classes (N= 34 subclusters and 12 cell classes). FDR was calculated for subclusters and cell classes separately. **e,** qPCR of pluripotent, primed, and naive markers correlates to cell class and subcluster proportions at day 84 (12/92 variables, 12/54 differentiations, 3 genes each for primed, naive and undifferentiated state, one gene each for three germ layers assessed). Technical correlates surviving FDR < 0.1 when correcting over all technical variables and genes tested are shown. No correlations at day 14 survived that threshold. Example relationships are highlighted: **f,** day 14 sc_2 are associated with CDH20 (endoderm) expression in iPSCs from the Taqman scorecard which is driven by one outlier, **g**, day 14 cortical hem proportions are associated with FGF lot number during reprogramming and **h,** day 84 PP/SP associated with SOX2 KLF4 $2^{-\Delta CT}$. KSR (knock out replacement serum) and L-glutamine refer to lot numbers for those reagents. Split_ratio_Set_1 refers to passaging the iPSCs after thawing at different ratios.

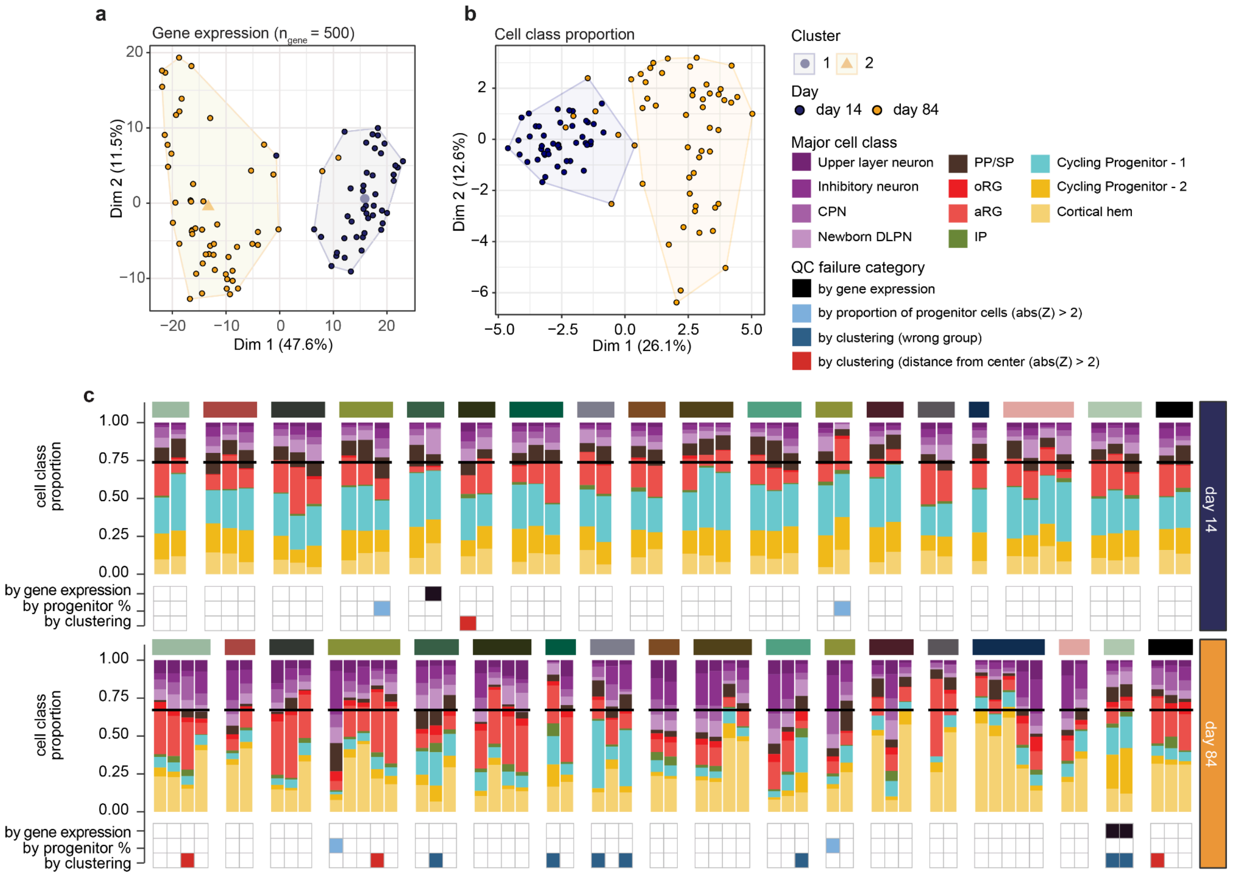


### Supplementary Figure 5| Reproducibility of cell type proportions and exclusion of poor differentiations

Clones categorized in the wrong cluster by either bulk gene expression (500 most variable autosomal genes) (**a**) or cell class proportion (**b**) were excluded from the analysis. We also removed clones if the proportions of neuronal cells (Maturing neuron, Inhibitory neuron, CPN, Newborn DLPN, and PP/SP) were outliers (abs(Z)>2) (**c**). In **c**, these QC metrics and the major cell class proportion for each sample are summarized. The horizontal line indicates the mean proportion of neuronal cells.

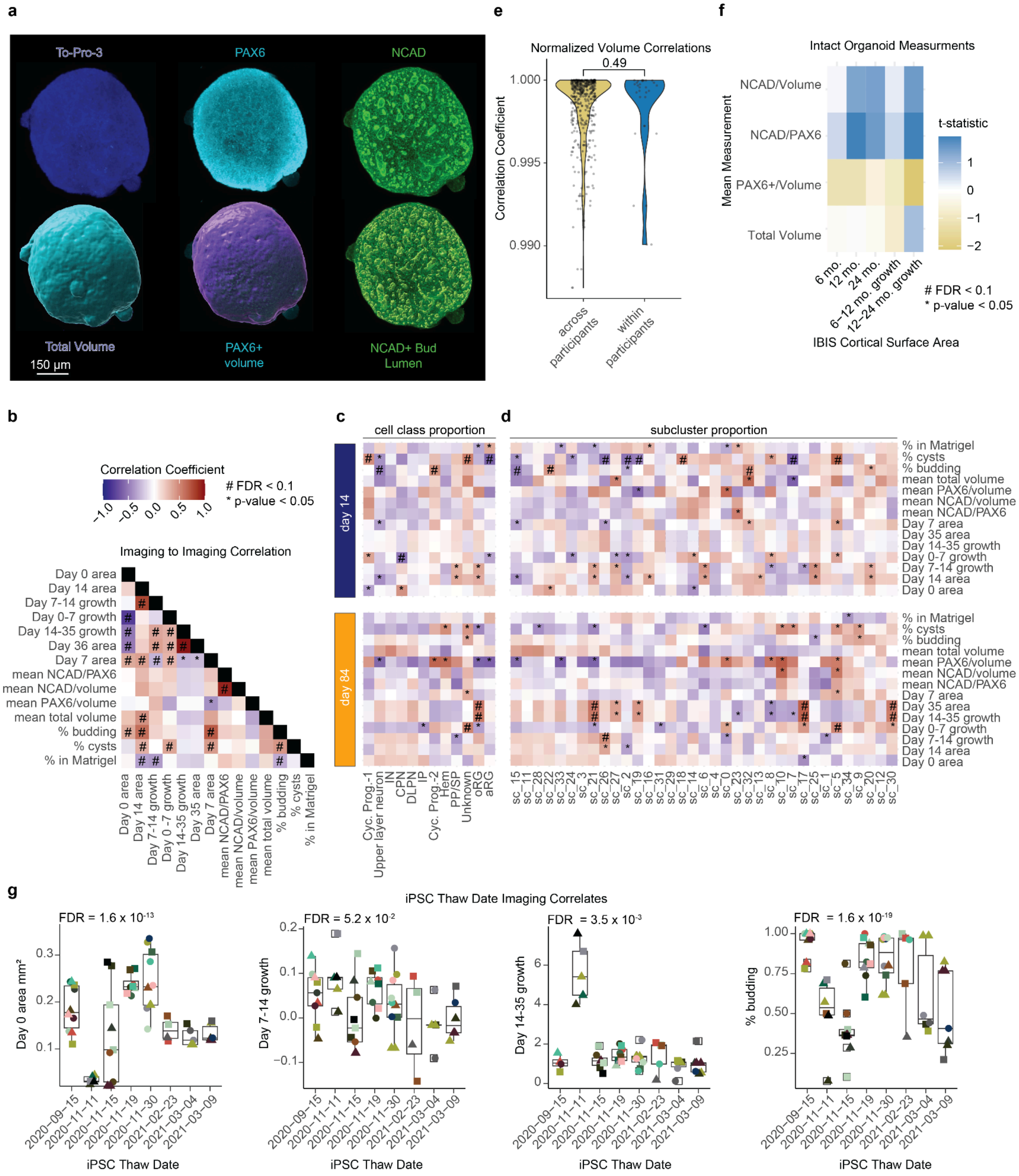


Supplementary Figure 6| Intact organoid imaging and technical correlates

**a.** Representative rendering of an immunolabeled intact day 14 organoid (top row) and volumetric analysis via segmentation in Imaris (bottom row). **b.** Pearson's correlations within hCO morphology, cross-sectional area and intact immunolabeling measurements (N = 14). **c**. Pearson's correlations to cell classes at day 14 and day 84 to hCO morphology, cross-sectional area and intact immunolabeling measurements. FDR was calculated separately for day 14 and day 84 (N = 14 imaging-based assays, N = 12 cell classes). **d**, Pearson's correlations to subclusters at day 14 and day 84 to hCO morphology, cross-sectional area and intact immunolabeling measurements. FDR was calculated separately for day 14 and day 84 (N = 14 imaging-based assays, N = 34 subclusters). **e.** Pearson correlation coefficients across and within participants for all measurements from intact immunolabeling. **f.** Volumetric measurement correlations to IBIS measurements. FDR correction was applied for each volumetric measurement to all MRI measurements. **g**, hCO cross-sectional area measurements and morphology that are influenced by the date when iPSC were thawed. Thaw date did not have a significant effect on day 7-14 growth rate.

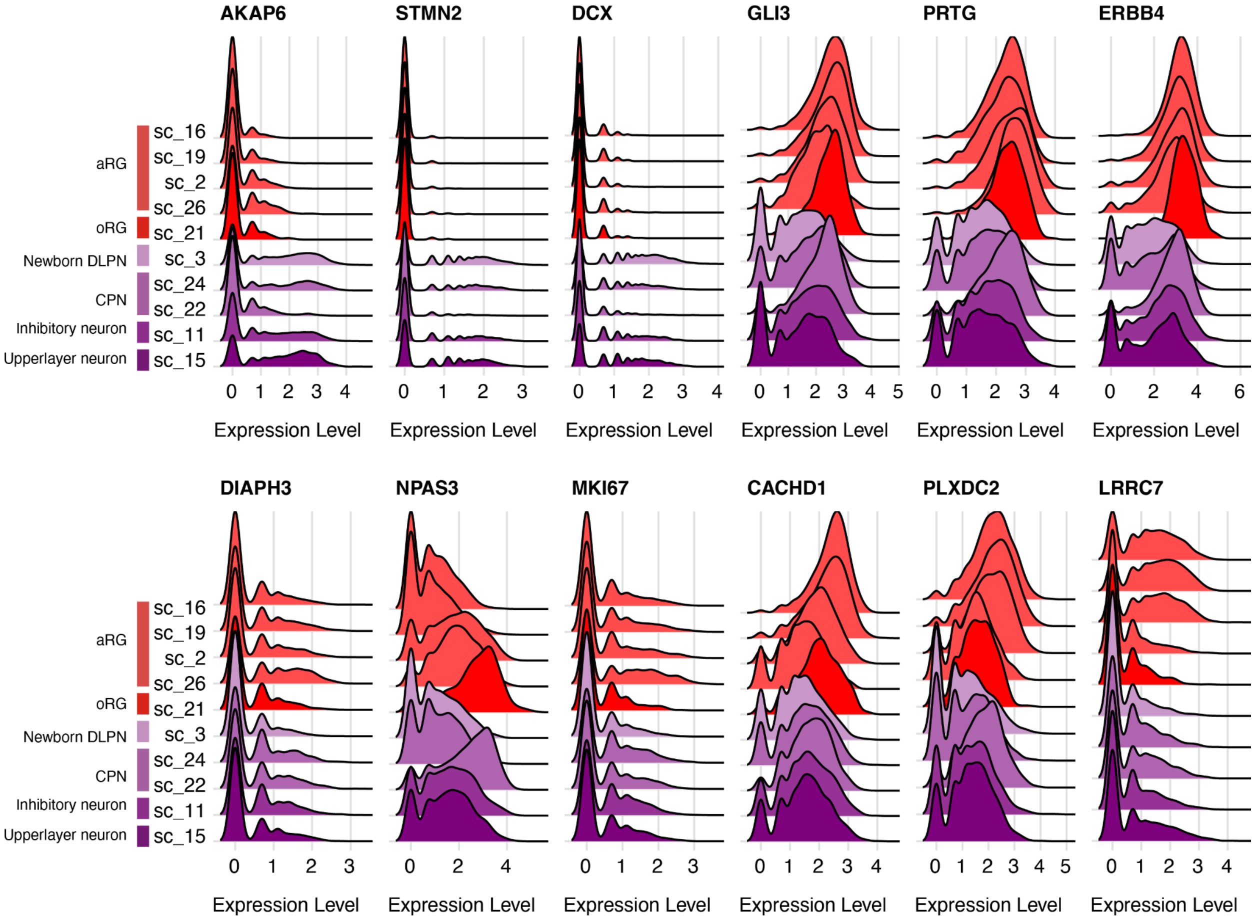


### Supplementary Figure 7| Top variable genes contributing pseudotime trajectory

Ridgepolot shows the gene expression in each subcluster colored by major cell class.

### Supplementary Tables

#### Supplementary Table 1| Participant included in each experiment

(Table.S1.DCCIDinExperiment)

#### Supplementary Table 2| Marker genes for scRNAseq subclusters and cell classes

(Table.S2.MarkerGenes.FDR01.xlsx)

#### Supplementary Table 3| Technical correlates to cell classes and subclusters

(Table.S3.TechnicaltoCellTypeProportions.xlsx)

#### Supplementary Table 4| Cell type and subcluster proportions to MRI measurements

(Table.S4.MRItoCellType.xlsx)

#### Supplementary Table 5| Technical correlates to hCO area and immunolabeling

(Table.S5.ImagingtoTechnical.xlsx)

#### Supplementary Table 6| Genes correlated to MRI measurements

(Table.S6.DEG.xlsx)

#### Supplementary Table 7| Enriched pathway in DEG

(Table.S7.Pathway.Enrichment.xlsx)
